## Supplemental Figures for "Interchromosomal segmental duplication drives translocation and loss of *P. falciparum* histidine-rich protein 3"

### SUPPORTING INFORMATION

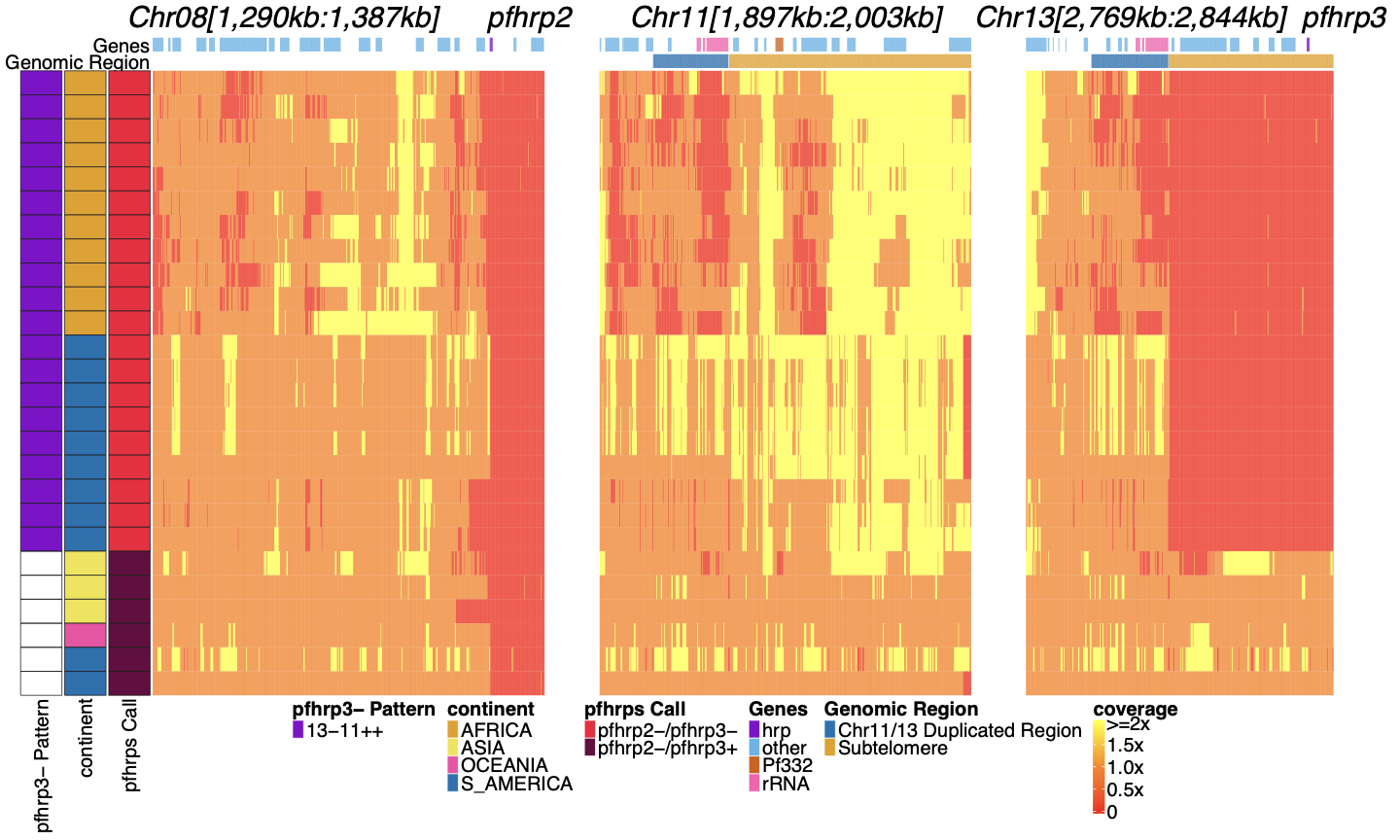

**Supplemental Figure 1****: Genome coverage of isolates with evidence of *pfhrp2* deletion.**

Sequence coverage heatmap of chromosomes 8 (1,290,239 - 1,387,982 bp), 11 (1,897,151 - 2,003,328 bp), 13 (2,769,916 - 2,844,785 bp). Displaying the 26 parasites out of the 19,313 samples with signs of possible *pfhrp2* deletions. Each row is a parasite. The top annotation along chromosomes depicts the location of genes, and the second row delineates the duplicated region (dark blue) and subtelomere region (orange). The left parasite annotation includes the deletion pattern, continent of origin, and *pfhrp2/3* deletion calls. The 20 parasites with evidence of HRP3 deletion were only found within South America (6 in Peru, 3 in Brazil) and Africa (11 in Ethiopia) and had evidence of the 13^-^11^++^ deletion HRP3 deletion pattern. Of the 6 parasites without HRP3 deletion (marked as white in *pfhrp3*- Pattern column for having no *pfhrp3* deletion), 2 were from South America, 3 from Asia (1 being lab isolate DD2), and 1 from Oceania (lab isolate D10).

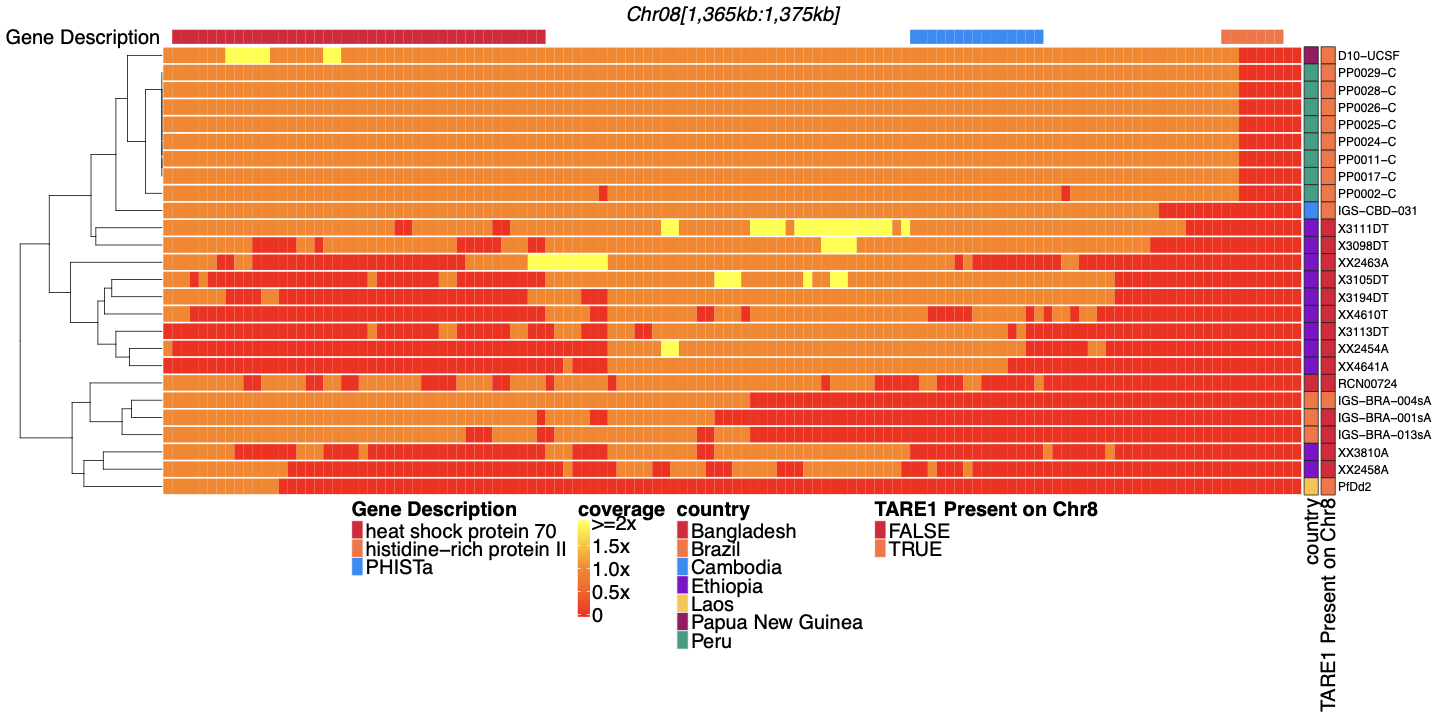

**Supplemental Figure 2**: **Coverage of sub-telomeric region of chromosome 8 before *pfhrp2* of parasites with *pfhrp2* deletion.**

Heatmap coverage normalized to genomic coverage of the sub-telomeric region of chromosome 8 (spanning 1,365,360-1,375,435 bp, 10,075bp in length) for the 26 parasites with *pfhrp2* genomic deletion. Each row is a parasite, and each column is a genomic location. The top annotates which gene the region falls within. The right side annotation shows the country of origin and which parasites have evidence of TARE1 at the location where genomic coverage drops to zero within this region. Most parasites without evidence of TARE1 or other genomic rearrangement are sWGA parasites and may lack the coverage to detect such events.

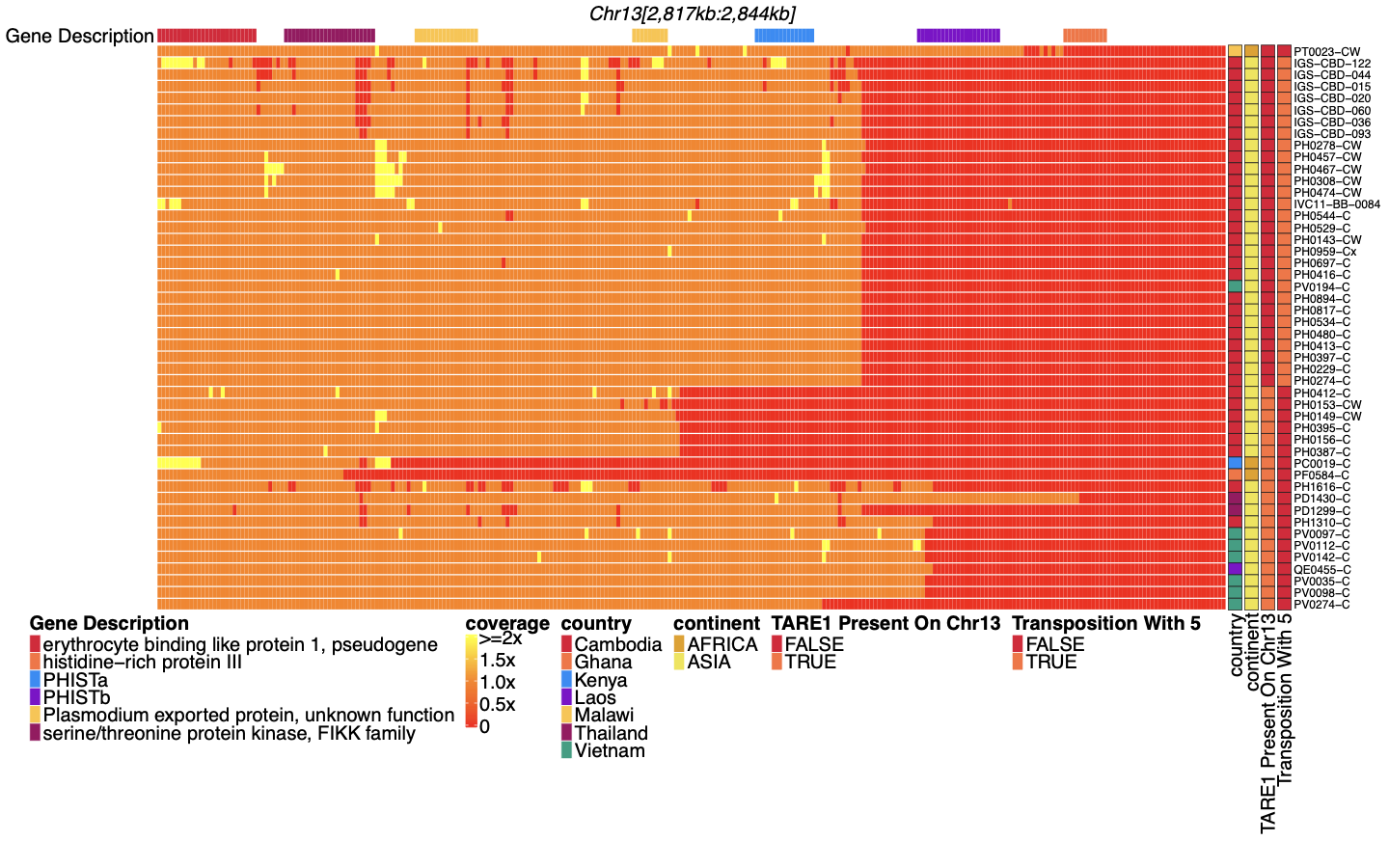

**Supplemental Figure 3**: **Coverage of chromosome 13 for parasites with** ***pfhrp3* deletion pattern 13^-^TARE1**.

Heatmap coverage normalized to genomic coverage of the sub-telomeric region of chromosome 13 (spanning 2,817,793 - 2,844,785 bp, 26,992bp in length) for the 48 parasites with *pfhrp3* deletions not associated with pattern 13^-^11^++^. Each row is a parasite, and each column is a genomic location. The top annotates which gene the region falls within. The right side annotation shows which parasites have evidence of TARE1 at the location where genomic coverage drops to zero within this region (n=19 **pattern 13^-^TARE**) and which parasites have evidence of genomic rearrangement with chromosome 5 from discordant paired-end reads, which results in duplication of *pfmdr1* (n=28 **pattern 13^-^5^++^**). The top parasite lacks evidence of either deletion type. The next 28 parasites have evidence of rearrangement with chromosome 5 with discordant reads with mates mapping to chromosome 13 and other mates mapping to chromosome 5. The next 19 parasites have evidence of TARE1 contiguous with chromosome 13 sequence on various locations consistent with breakage and telomere healing. These breaks occur on chromosome 13 at 2,836,793 (n=8), 2,830,793 (n=4), 2,829,793 (n=2), 2,821,793 (n=1), 2,822,793 (n=1), 2,833,793 (n=1), 2,834,793 (n=1), 2,836,793 (n=1), and 2,840,793 (n=1).

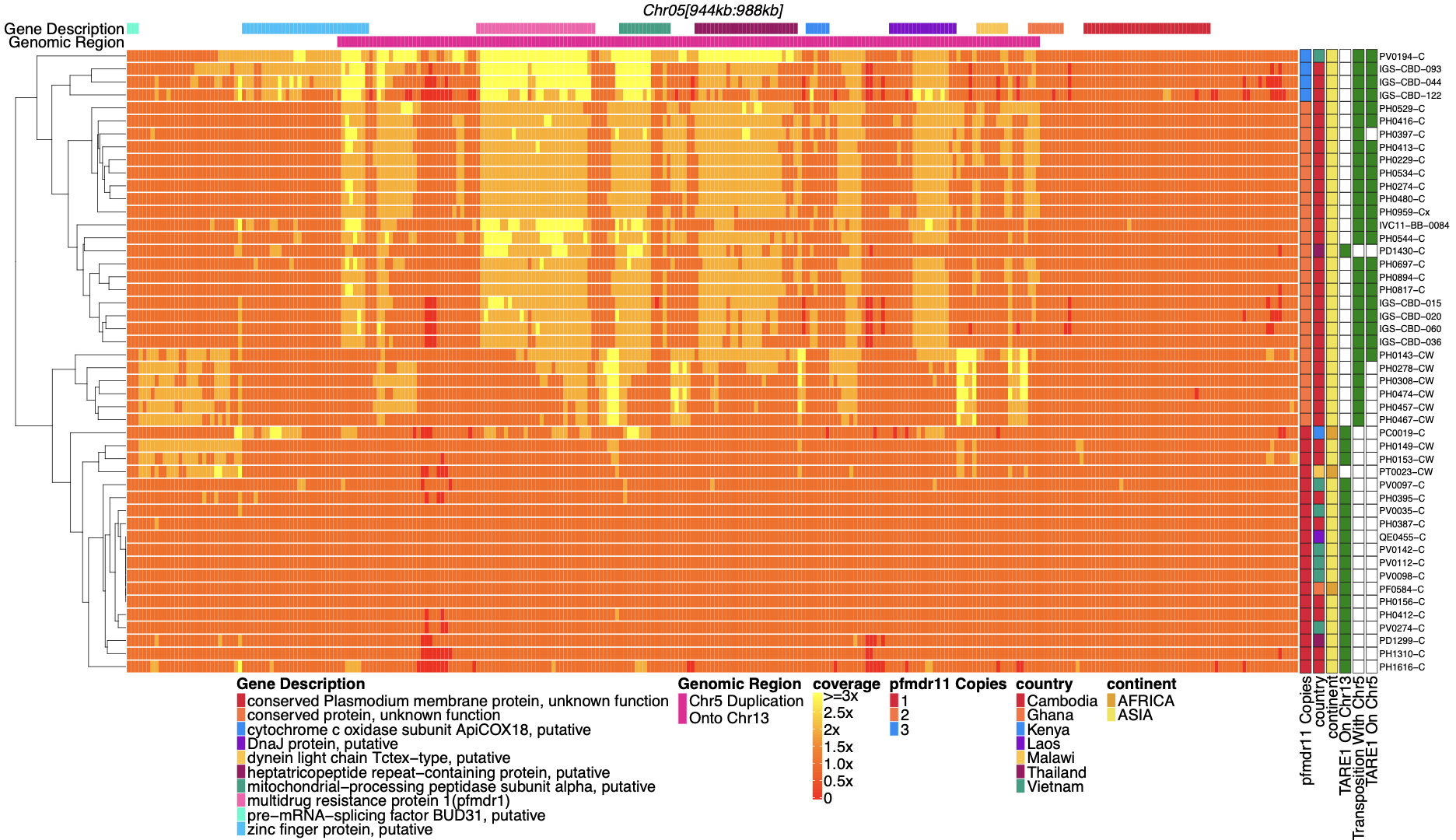

**Supplemental Figure 4**: **Coverage of chromosome 5 for parasites with** ***pfhrp3* deletion pattern 13^-^TARE1 and 13^-^5^++^**.

Heatmap coverage normalized to genomic coverage of a region of chromosome 5 (spanning 944,534 - 988,747 bp, 44,213bp in length) for parasites with *pfhrp3* deletion pattern 13^-^TARE1. Each row is a parasite, and each column is a genomic location. The top annotates which gene the region falls within. The right side annotates the country of origin of the parasite. The 3rd annotation bar on the right side shows which parasites have TARE1 detected on chromosome 13; parasites with green have TARE1 detected on chromosome 13 that are clustered on the bottom of the graph and have normal coverage across this region of chromosome 5, while the parasites on top have evidence of re-arrangement between chromosome 13 (position 2,835,587) and chromosome 5 (position 979,203) and show increased coverage across chromosome 5 up to the point where TARE1 sequence is detected on the reverse strand. The second annotation hot-pink bar from the top indicates the duplicated region. The beginning of this bar is where the TARE1 sequence on the reverse strand is detected. The above would be consistent with a genomic rearrangement between chromosome 13 at position 2,835,587 and chromosome 5 at position 979,203, which results in the deletion of chromosome 13 from 2,835,587 onwards and the duplication of a 26KB region of the reverse strand of chromosome 5 from position 979,203 to 952,668 resulting in the duplication of *pfmdr1* and the deletion of *pfhrp3*. The parasites with transposition evidence between 13 and 5 but no TARE1 detected on 5 were all sWGA parasites, which likely limited the ability to detect the TARE1 sequence. The top 4 parasites appear coverage-wise to have 3 copies of *pfmdr1*. The top 4 parasites, in addition to the likely copy on chromosome 13, have a tandem duplication on chromosome 5. There were 2 different tandem duplications detected. The 1st parasite is tandemly duplicated from a monomeric stretch of Ax19 at 947,967 and Ax18 at 970,043 and is from Laos. The 2nd, 3rd, and 4th parasites, all from Cambodia, are tandemly duplicated from Ax22 at chromosome 5 946,684 and Ax36 at 964,504. All parasites with evidence for 13^-^5^++^ have only wild-type *pfmdr1* or are a mix of wild-type and 184F. The four parasites with three copies of *pfmdr1* have 2x coverage of 184F and 1x coverage of wild-type *pfmdr1*, which is consistent with the duplicated *pfmdr1* on chromosome 13 being wild-type *pfmdr1*. All parasites with this deletion pattern are found within Asia.

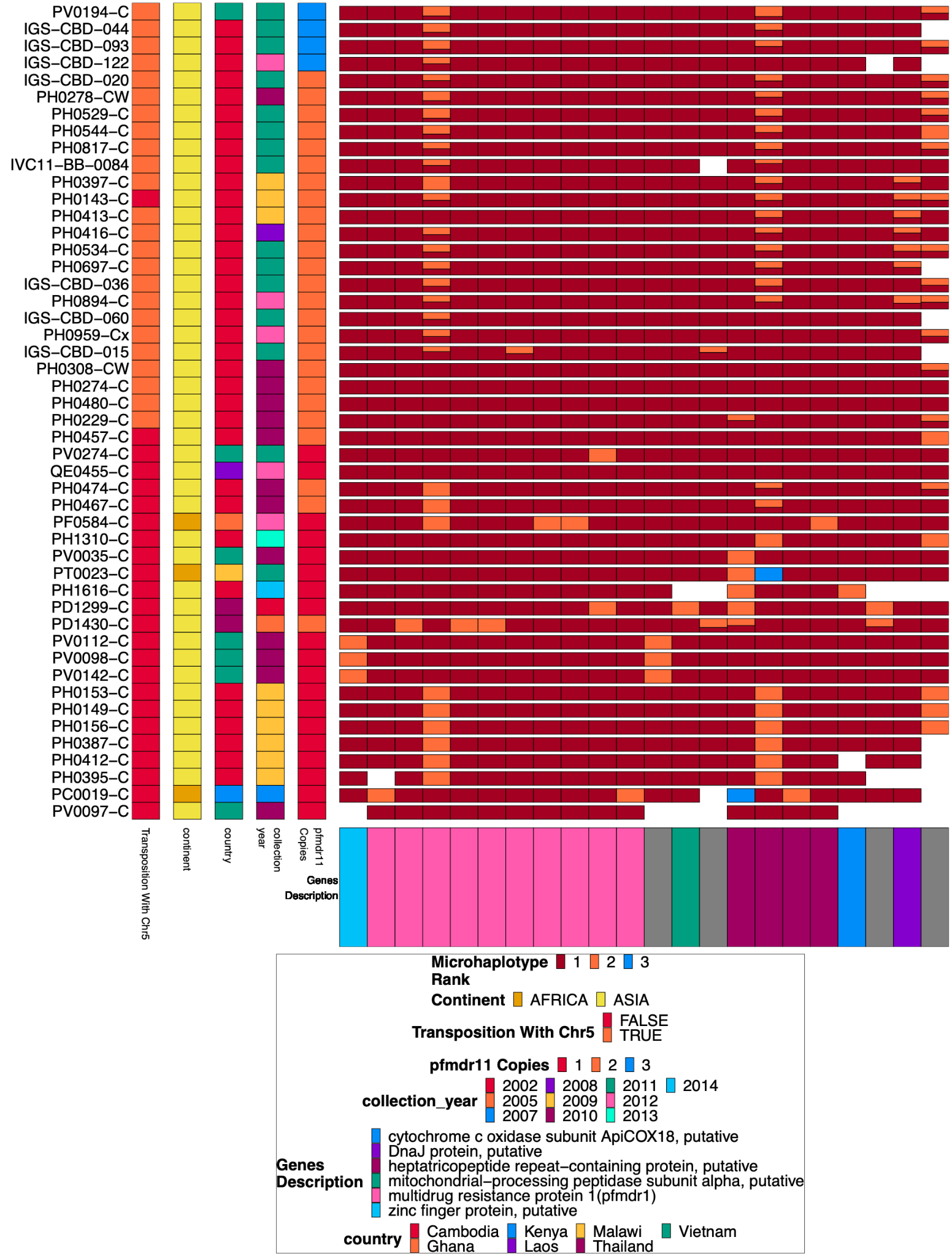

**Supplemental Figure 5**: **Chromosome 5 duplicated region microhaplotypes.**

The 22 microhaplotype regions with variation across the duplicated portion of chromosome 5 spanning from 952,668 to 979,203 for the 48 isolates with chromosome 13 deletion without chromosome 11 duplication. Each row is an isolate. In each column, the isolate is typed by microhaplotype (colored by the prevalence of each microhaplotype, with 1=red being most prevalent, 2=orange second most prevalent, and 3=blue least prevalent). This color coding system is specific to each column, and the same color across columns does not indicate the same haplotype, just the prevalence in the population for that column. Associated metadata for each isolate can be seen on the left after the isolate’s name. Most (n=28) of these show evidence of complex recombination with chromosome 5 at 952,668 and 2,835,587 on chromosome 13, resulting in the deletion of *pfhrp3* and the duplication of *pfmdr1*. Only 6 isolates with MDR duplication have no variation within *pfmdr1*. The other isolates have a wild type Y184 (yellow) on one copy and 184F (red) on the other copy.

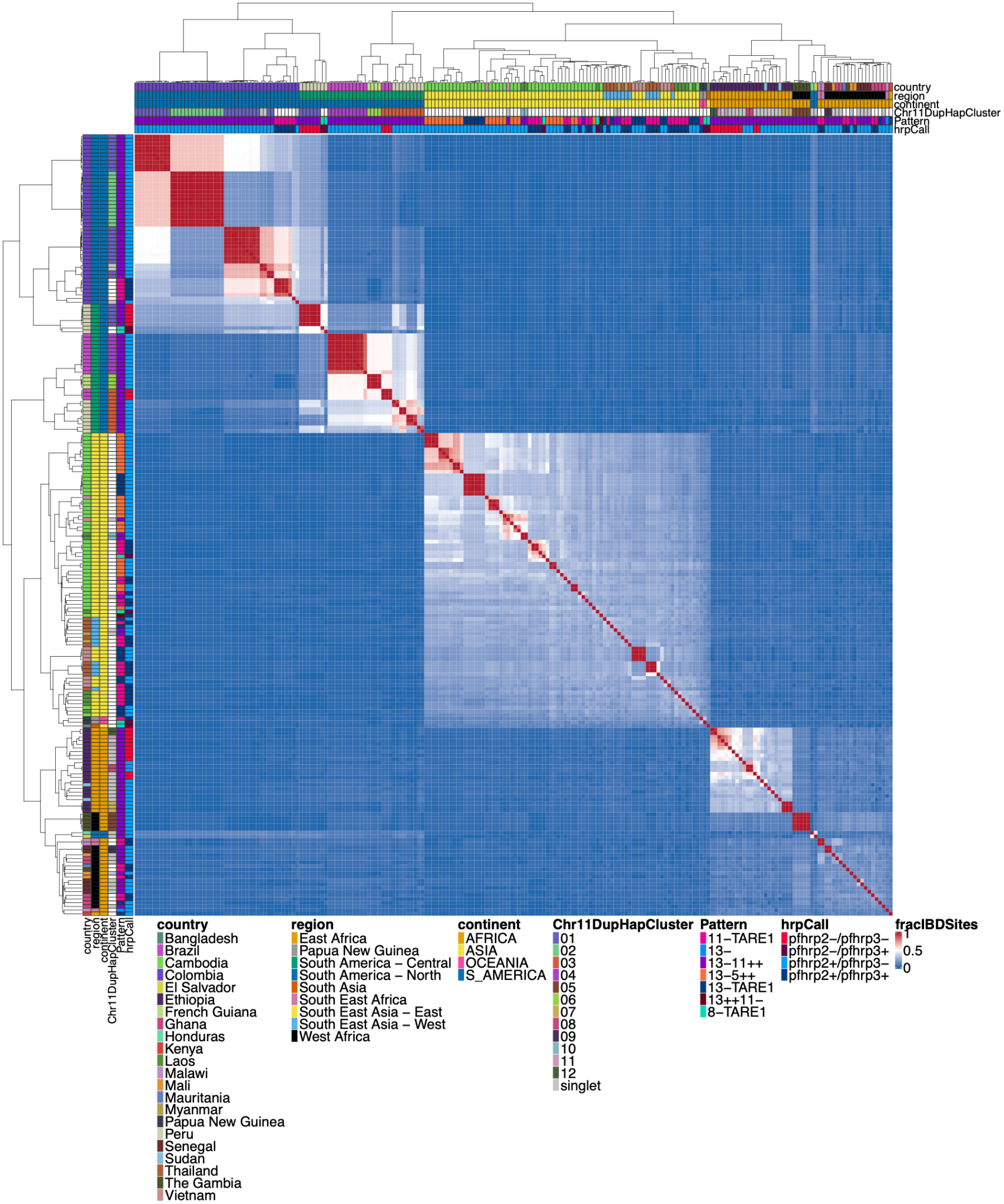

**Supplemental Figure 6**: **Whole genome** **IBD between** **all parasites with a genomic deletion**

Whole genomic IBD was calculated between all parasites within this study with a genomic deletion (on chromosome 8, 11, or 13) using hmmIBD and whole genome biallelic SNPs. The fraction of sites in IBD between parasites was plotted above. The top and left side annotations are the same and contain the parasite's country, region, and continent of origin. The annotation also has the deletion calls for *pfhrp2/3*, the deletion Pattern, and which chromosome 11 haplotype cluster the parasite belongs to. The parasites cluster strongly by continent and by subregion/country. The two South American parasites clustered within the African samples are lab isolates HB3 and Santa-Luca-Salvador-1.

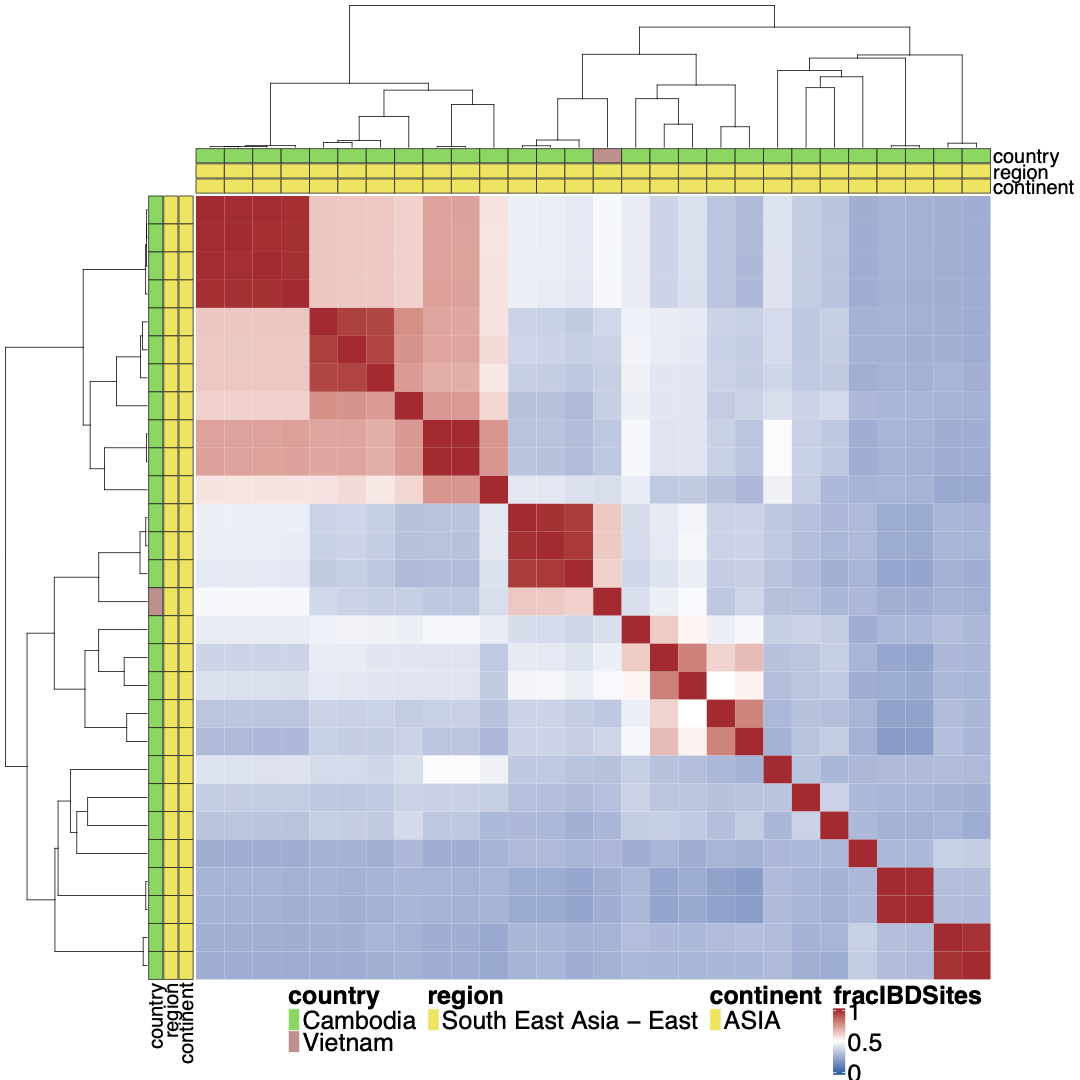

**Supplemental Figure 7**: **Whole genome** **IBD between** ***pfhrp3* deletion pattern 13^-^5^++^ parasites**

Whole genome IBD was calculated between *pfhrp3* deletion pattern 13^-^5^++^ parasites using hmmIBD using whole genome biallelic SNPs. The fraction of sites in IBD between parasites was plotted above. The top and left side annotations are the same and contain the parasite's country, region, and continent of origin. Haplotypes do not appear like a clonal expansion of a parasite containing the 13-5 hybrid chromosome, and either multiple different events occurred to create this hybrid translocation, or it occurred once, and the hybrid 13-5 chromosome has been passed onto prodigies which have continued to undergo sexual recombination decreasing their whole genome IBD.

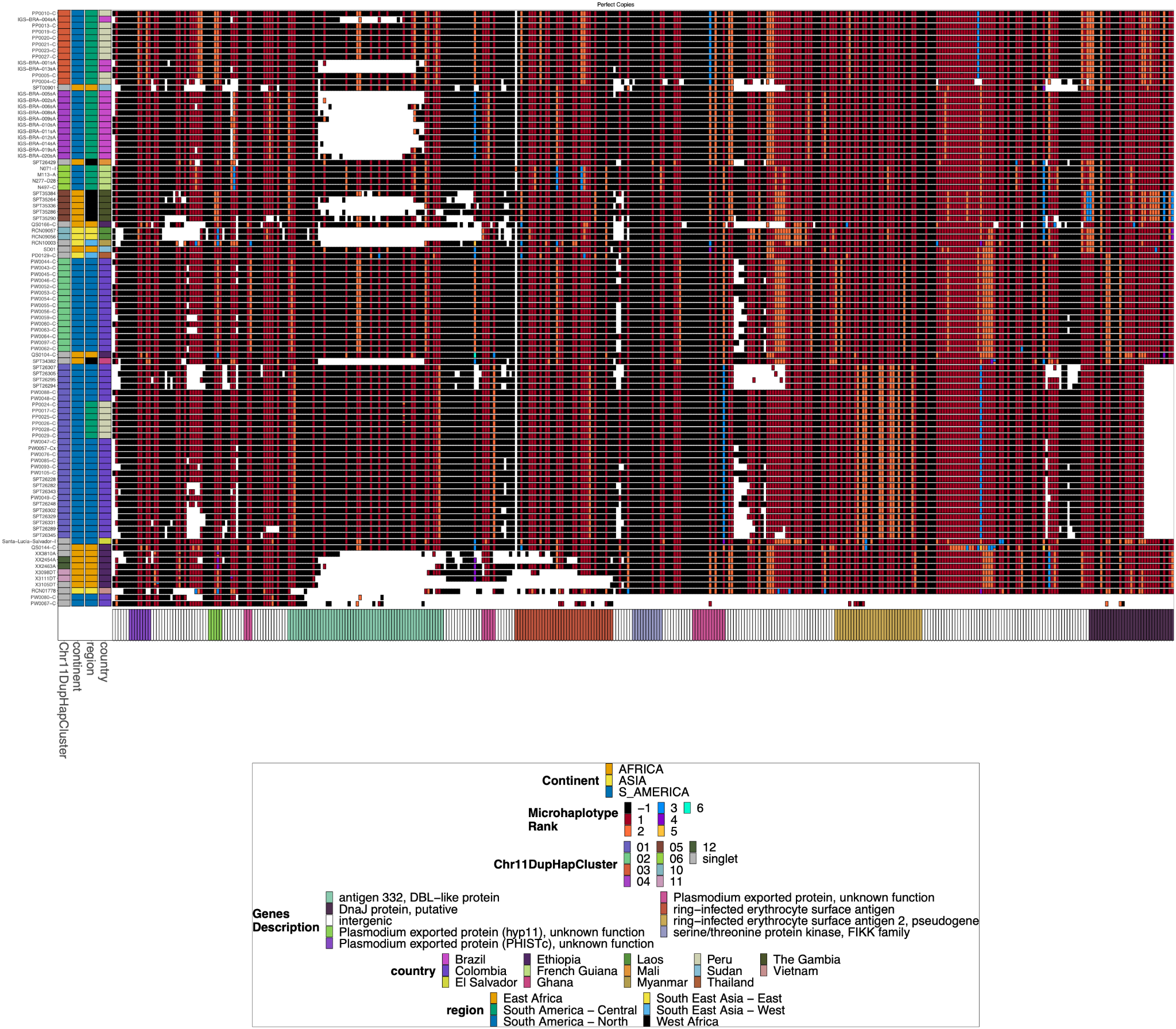

**Supplemental Figure 8**: **Chromosome 11 Duplicated Segment *pfhrp3* deletion Pattern 13^-^11^++^ parasites with perfect copies.**

Subset of the parasites from **Supplemental Figure 11** for the parasites that have a perfect duplication of the chromosome 11 segment. There are clear divergent haplotypes (9 can be found in multiple samples, with the largest group being found in 28 samples) for the perfect duplications, consistent with the duplication event occurring multiple times and does not stem from a single event from which all parasites are descended.

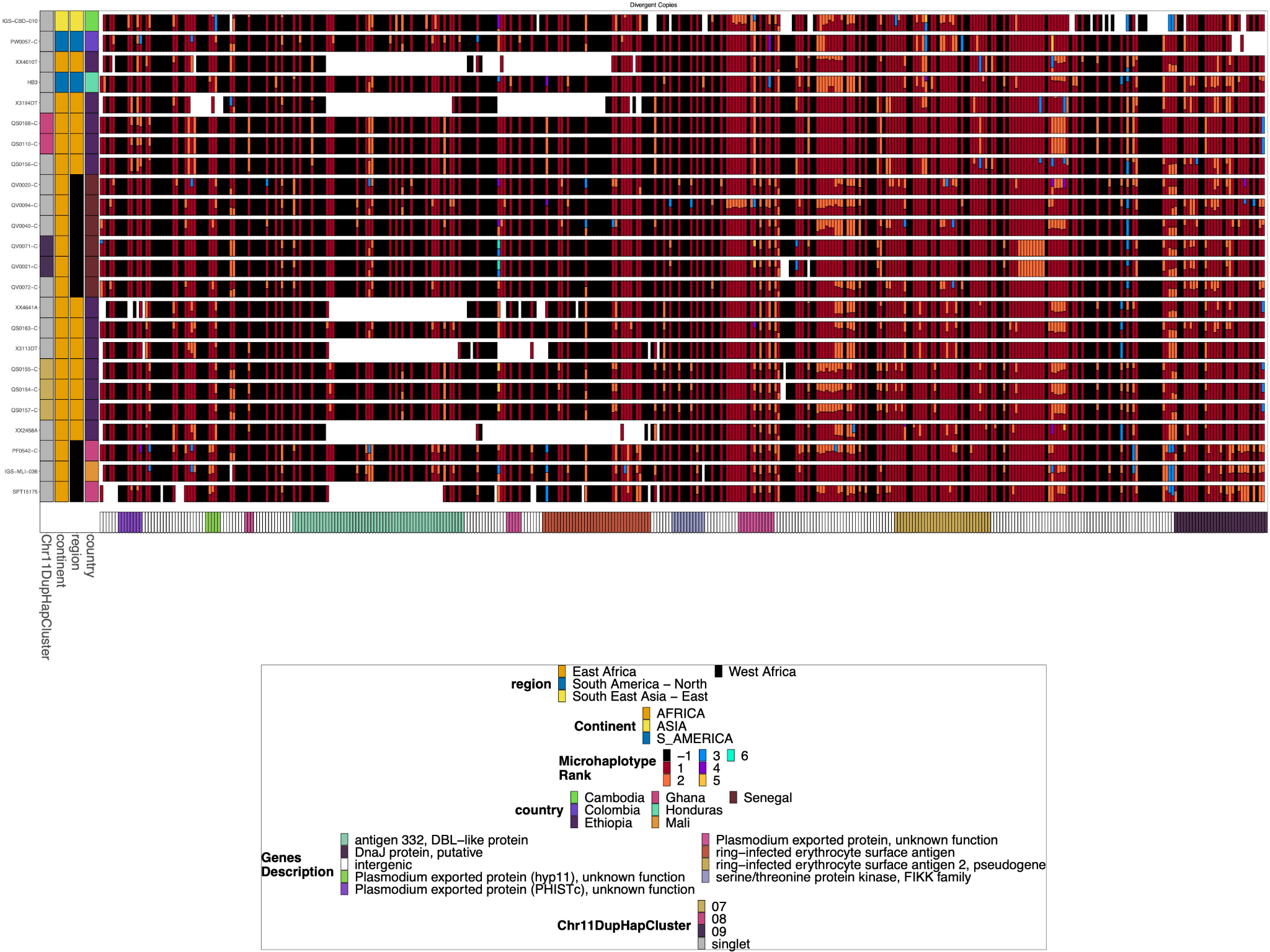

**Supplemental Figure 9**: **Chromosome 11 Duplicated Segment *pfhrp3* deletion Pattern 13^-^11^+^**^+^ **parasites with divergent chromosome 11 copies.**

Subset of the parasites from **Supplemental Figure 11** for the parasites that have divergent duplicates of the chromosome 11 segment. Several parasites have divergent chromosome 11 segments, but they share the same exact divergent copies with other parasites, which would be consistent with the coinheritance of the two divergent copies simultaneously. This could be consistent with parasites inheriting from previous duplication events involving divergent copies or meiotic recombination between parasites with two separate duplication events of disparate chromosome 11 segments, inheriting one chromosome 11 segment on chromosome 13 from parent 1 and a different chromosome 11 segment on chromosome 11 from parent 2.

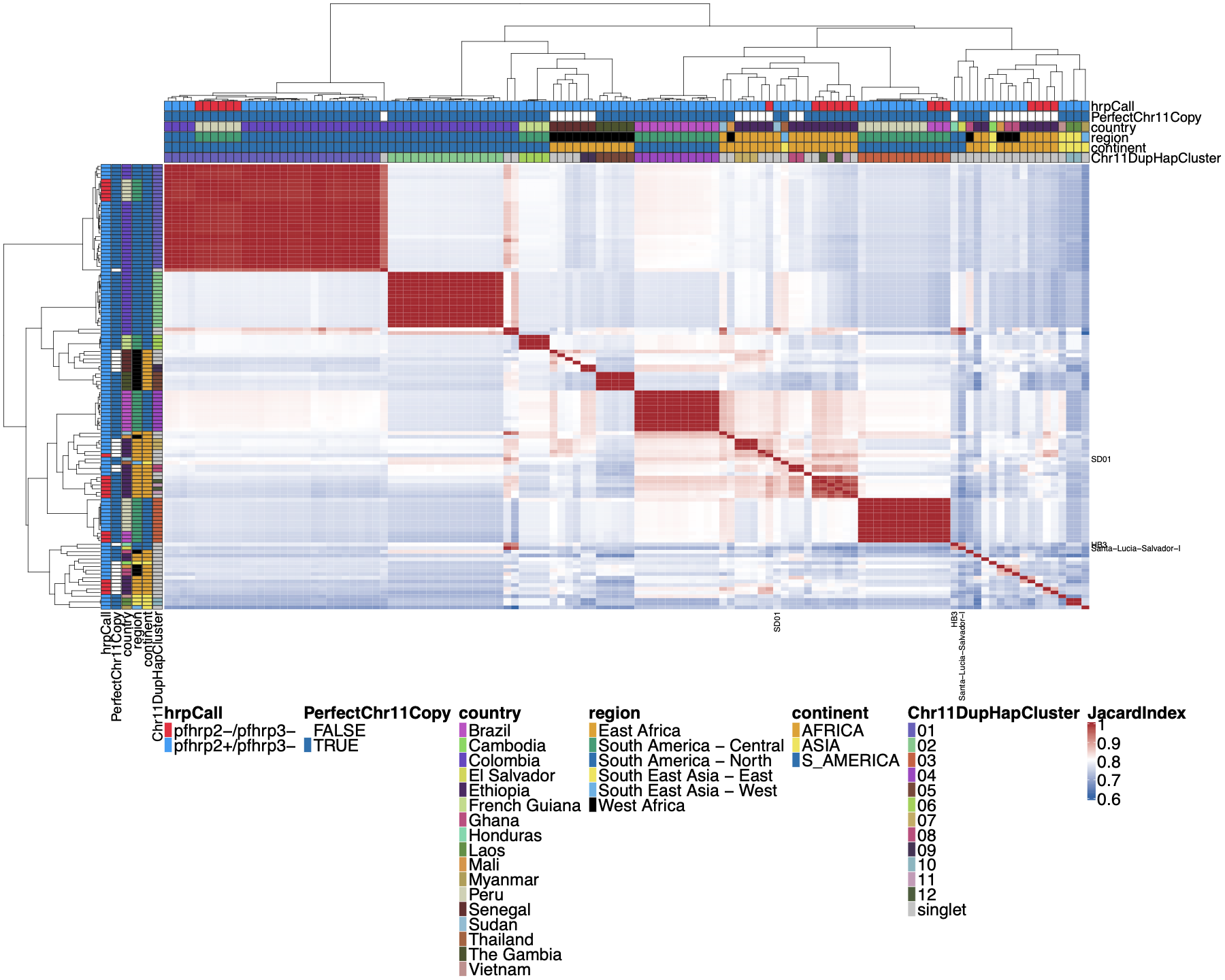

**Supplemental Figure 10**: **Jaccard similarity between parasites for chromosome 11 duplicated segment for *pfhrp3* deletion pattern 13^-^11^++^ parasites.**

This is an all-by-all distance matrix showing Jaccard similarity for the duplicated chromosome 11 segments between all parasites with pfhrp3 deletion pattern 13^-^11^++^. The parasites’ continent, region, and country are annotated on the sides of the heatmap, along with the *pfhrp2/3* deletion calls and whether the chromosome 11 duplicated segment shows an identical haplotype across the chromosome 11 duplicated segment. There are clearly several different haplotypes within the duplicated chromosome 11 segment, and there does not appear to be one specific haplotype associated with the duplication. The haplotypes found in more than 1 sample are clustered into “Chr11DupHapClusters” and colored in the annotation bar. Parasites group strongly by geographical location.

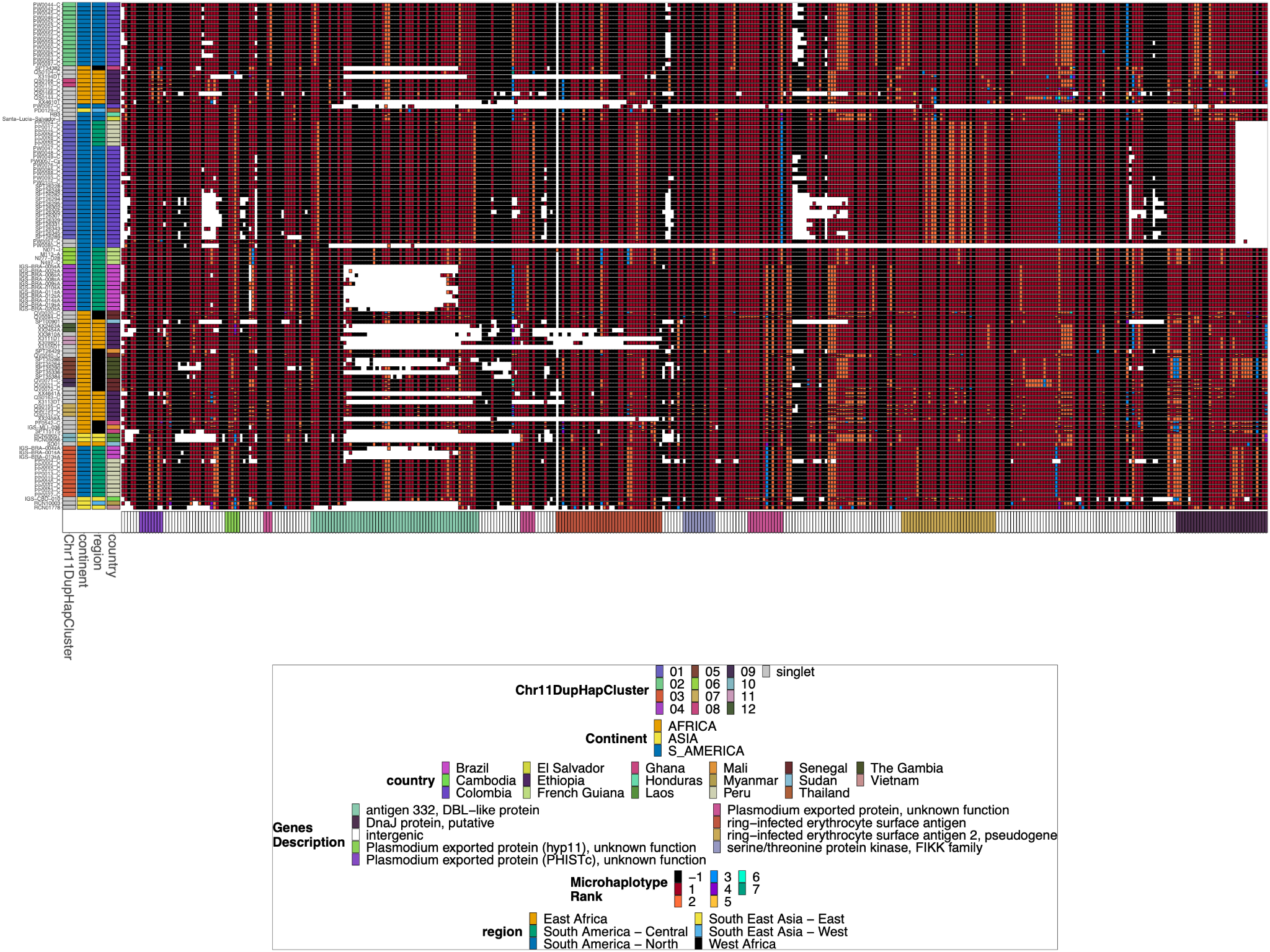

**Supplemental Figure 11****: Chromosome 11 Duplicated Segment *pfhrp3* deletion Pattern 13^-^11^++^ parasites.**

Plotted haplotype variation per sub-genomic regions across the duplicated chromosome 11 segment for the *pfhrp3* pattern 13^-^11^++^ parasites. Across the x-axis are the genomic regions in genomic order, and the genomic region genes are colored on the bottom bar. Each genomic area is slightly different in size, and there is genomic space between each region, so the plot is not on a genomic scale. The y-axis is each parasite with pattern 13^-^11^++^ of *pfhrp3* deletion where this segment of chromosome 11 is duplicated onto chromosome 13. The continent, region, and country are colored per parasite on the leftmost of the plot. Each column contains the haplotypes for that genomic region colored by the haplotype prevalence rank (more prevalent have a lower rank number, with most prevalent having rank 1) at that window/column. Colors are by frequency rank of the haplotypes (most prevalent haplotypes have rank 1 and colored red, 2nd most prevalent haplotypes are rank 2 and colored orange, and so forth. Shared colors between columns do not mean they are the same haplotype. If the column is black, there is no variation at that genomic window. If there is more than one variant for a parasite at a genomic location, the bar’s height is the relative within-parasite frequency of that haplotype for that parasite. The parasites are in the same order as the heatmap dendrogram in **Supplemental Figure 10**. There are clear, distinctive haplotypes for this duplicated region. The leftmost annotation bar shows which cluster (based on the sharing >99% identity in this area) the parasite belongs to, which is named “Chr11DupHapCluster”.

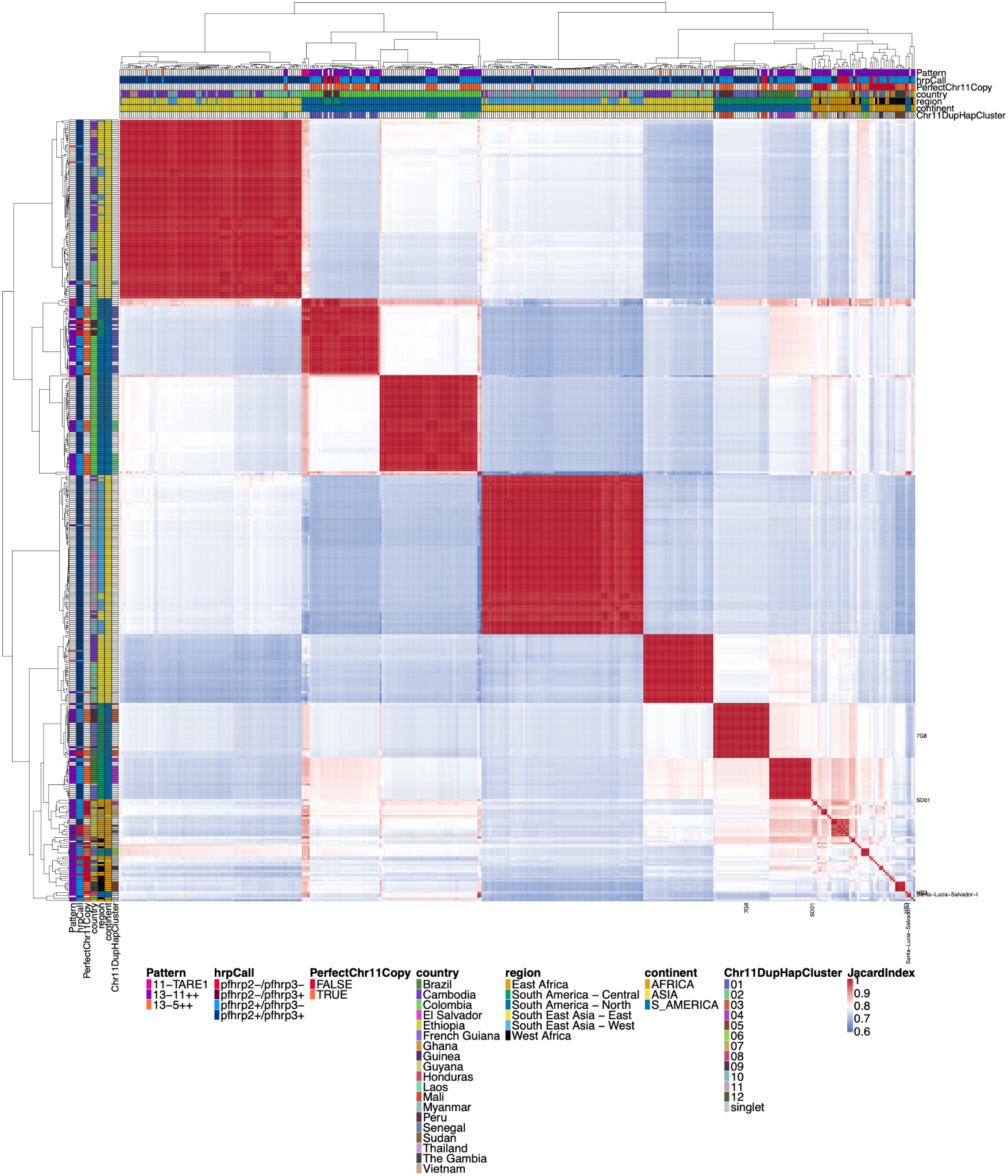

**Supplemental Figure 12**: **Jaccard similarity for chromosome 11 duplicated segment**

All parasites with micohaplotypes similar to the duplicated chromosome 11 microhaplotypes for the *pfhrp3* deletion Pattern 13^-^11^++^ parasites**.** While similar to **Supplemental Figure 10**, this all-by-all heatmap of Jaccard similarity includes all parasites with a similar chromosome segment to the parasites with pattern 13^-^11^++^ *pfhrp3* deletions. For the side and top annotations of parasites that do not have chromosome 11 duplication, there is a white bar for whether or not they have perfect chromosome 11 duplication. There are many parasites with closely related chromosome 11 segments to the duplicated chromosome 11 segments, indicating that the duplicated chromosome 11 segments also circulate within the population in strains with normal chromosome 11 and 13 arrangements.

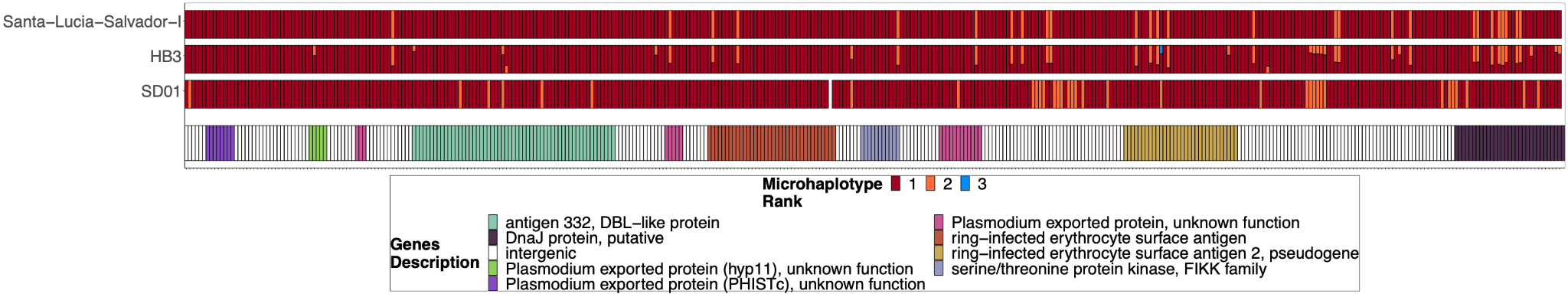

**Supplemental Figure 13****: Chromosome 11 Duplicated Segment coverage for *pfhrp3* deletion Pattern 13^-^11^++^ parasites SD01, HB3, and Salvador 1.**

Subset of the parasites from **Supplemental Figure 11** but for SD01 and HB3, which were sequenced in this paper, and for Santa-Luca-Salvador-1, another lab isolate that shows similar *pfhrp3* deletion pattern 13^-^11^++^. SD01 and Santa-Luca-Salvador-1 have perfect copies, while HB3 has divergent copies.

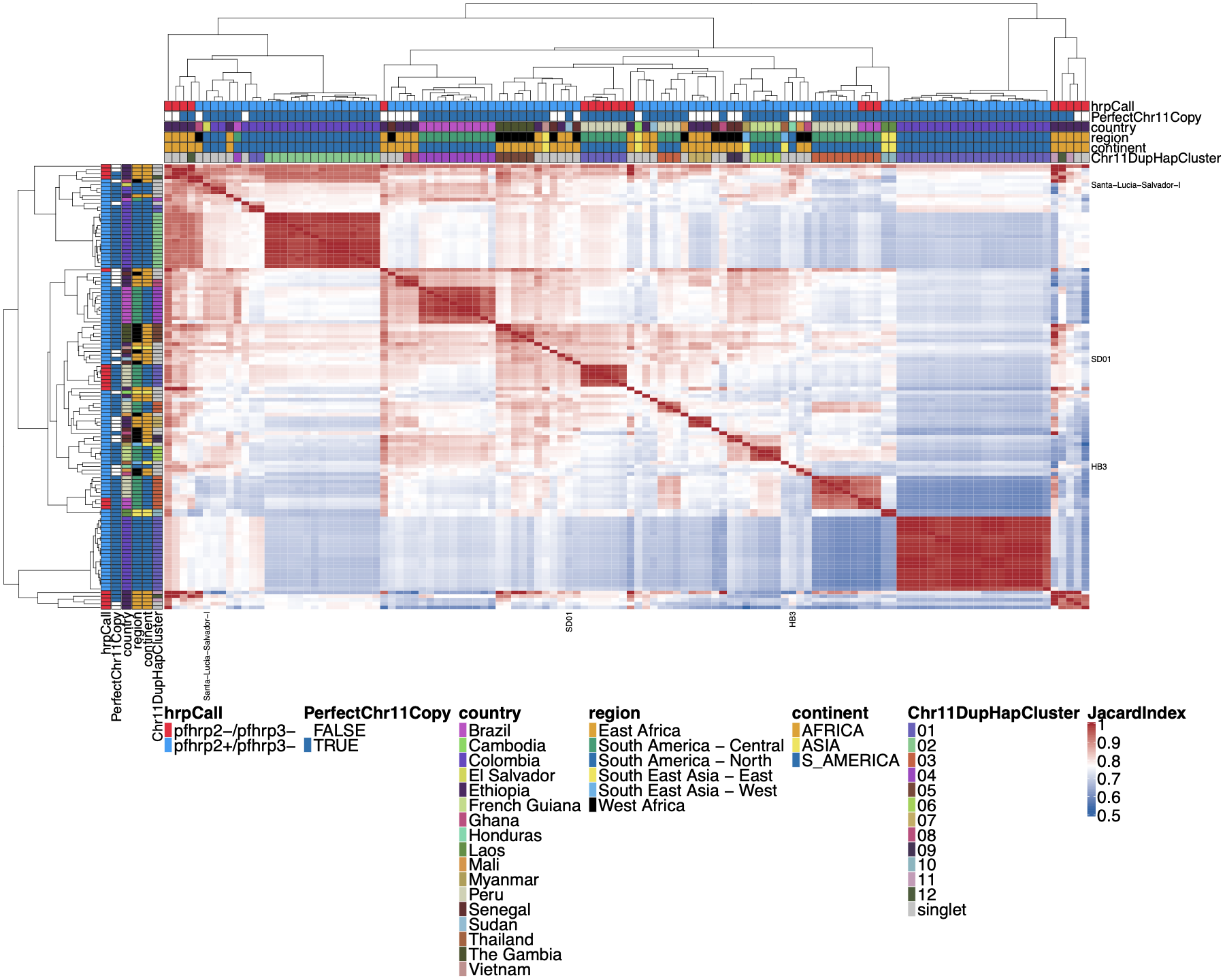

**Supplemental Figure 14****: Jaccard similarity for Chromosome 11/13 15.2kb duplicated region for *pfhrp3* deletion pattern 13^-^11^++^ parasites.**

An all-by-all distance matrix showing Jaccard similarity for the chromosome 11 and 13 duplicated region between all the parasites with *pfhrp3* deletion pattern 13^-^11^++^. The top triangle is identical to the bottom triangle. Parasites’ continent, region, and country are annotated on the sides of the heatmap, as well as the *pfhrp2/3* deletion calls and whether the chromosome 11 duplicated segment is a perfect copy. Sequences cluster loosely per geographic region, with similar sequences from the same country. However, parasites are not as strong separately by continent as for the duplicated chromosome 11 segment. Despite all parasites having duplicated chromosome 11 via this region, there are clearly different haplotype groups, consistent with multiple origins of this duplication event.

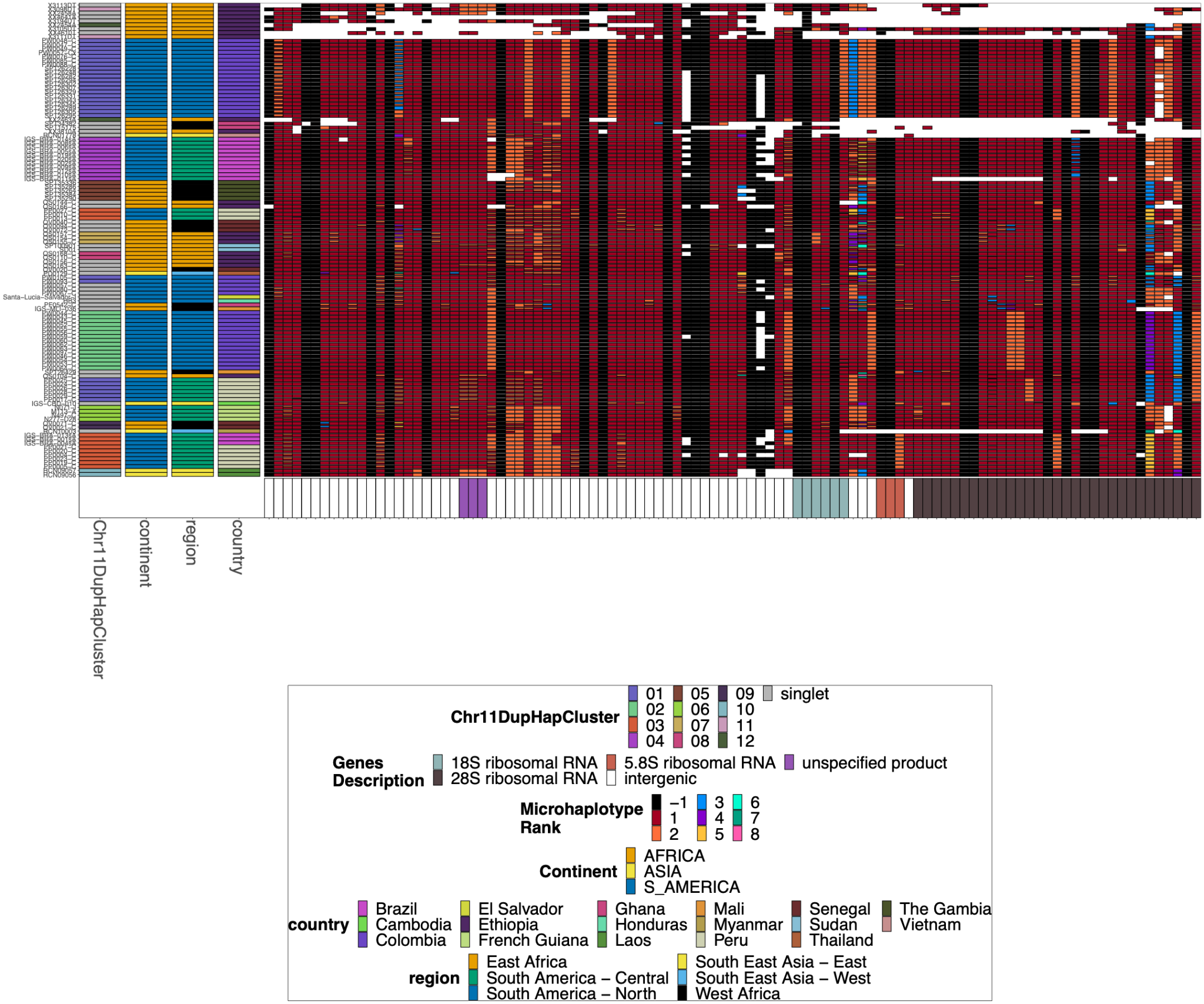

**Supplemental Figure 15**: **Chromosome 11/13 15.2kb duplicated region for *pfhrp3* deletion pattern 13^-^11^++^ parasites.**

Plotted microhaplotype variation across the region shared between all chromosomes 11 and 13. Across the x-axis are the genomic regions in genomic order, and the genomic region genes are colored on the bottom bar. The y-axis is each parasite with pattern 13^-^11^++^ of *pfhrp3* deletion where this segment of chromosome 11 is duplicated onto chromosome 13. The continent, region, and country are colored per parasite on the left most of the plot. Each column contains the haplotypes for that genomic region colored by the haplotype prevalence rank (more prevalent have a lower rank number, with most prevalent having rank 1) at that window/column. Colors are by frequency rank of the haplotypes (most prevalent haplotypes have rank 1 and colored red, 2nd most prevalent haplotypes are rank 2 and colored orange, and so forth. Shared colors between columns do not mean they are the same haplotype. If the column is black, there is no variation at that genomic window. If there is more than one variant for a parasite at a genomic location, the bar’s height is the relative within-parasite frequency of that microhaplotype for that parasite. Pattern 13^-^11^++^ is missing 46,323 bases from chromosome 13 (2,807,159 to 2,853,482) with a gain of 70,190 bases of chromosome 11 (1,933,138 to 2,003,328). Based on genomes that are assembled to the end of their telomeres(Otto et al., 2018a), an additional 17-84kb is deleted from the paralogous sub-telomeric region on chromosome 13, and an additional 15-87kb of the paralogous sub-telomeric region on chromosome 11 is duplicated.

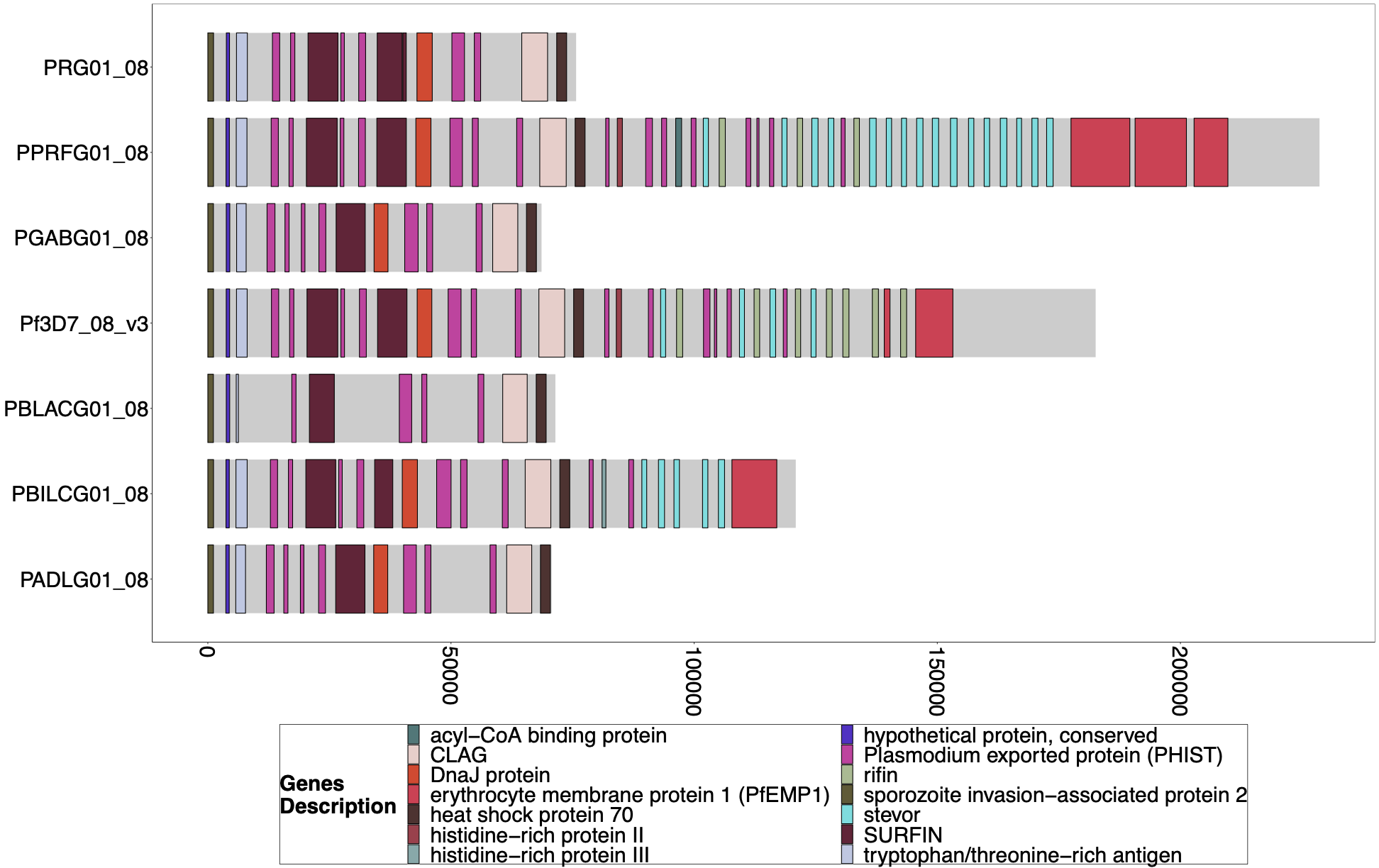

**Supplemental Figure 16****: Gene Annotations of Chromosome 8 of PacBio-assembled *P. Laverania* Genomes.**

Plots of the peri-telomere regions of chromosome 8 across all sequenced *Laverania* Genomes(Otto et al., 2018b). This region's assembly is incomplete for most strains, and only *Pf3d7* and the closest relative to falciparum, *PPRFG01*, contain *hrp2*, but they are in similar locations. The bottom x-axis shows the genomic distance from the starting point in each strain but there is significantly more core genome to the left of the starting point.

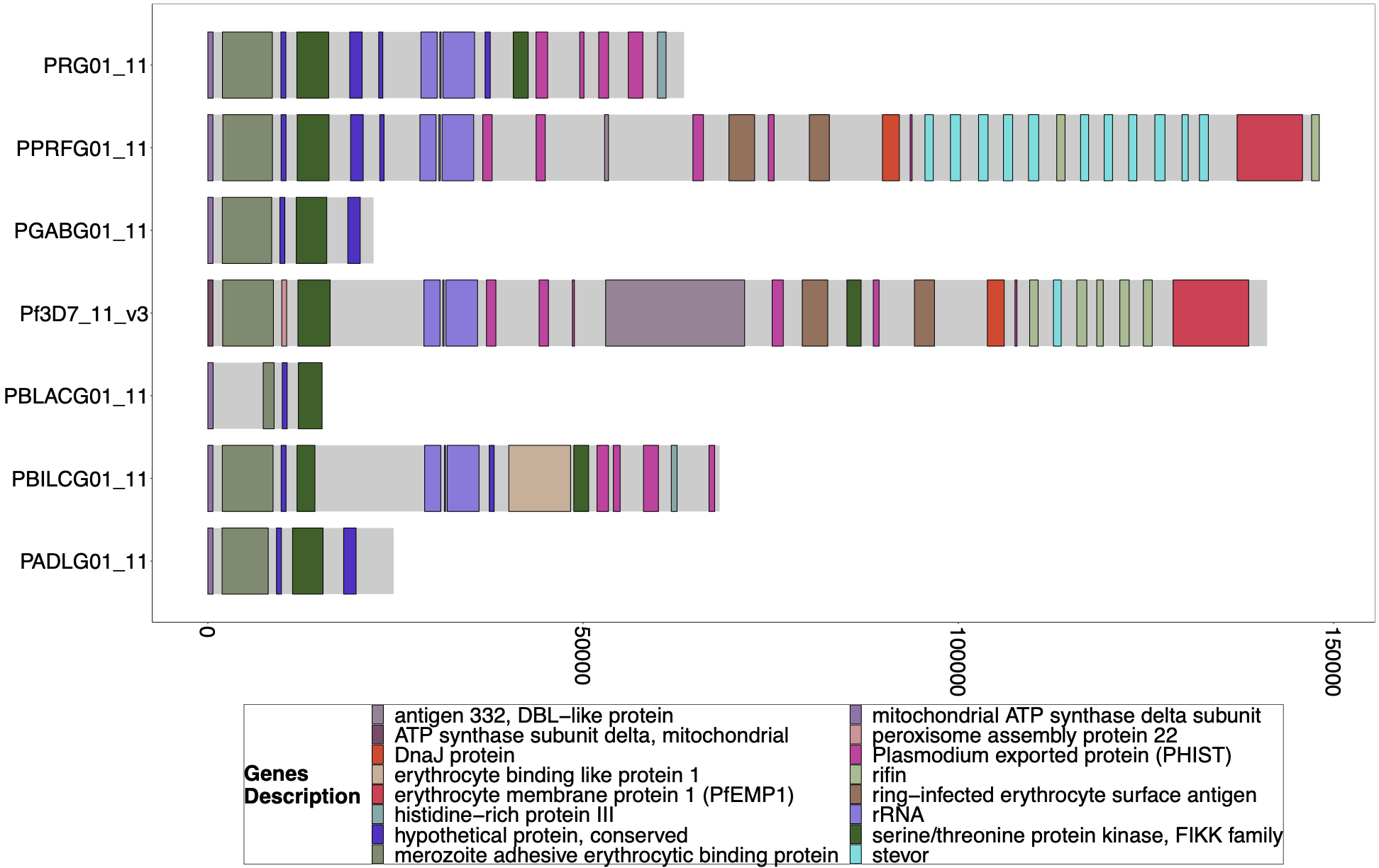

**Supplemental Figure 17****: Gene Annotations of Chromosome 11 of PacBio-assembled *P. Laverania* Genomes**

Plots of the peri-telomere regions of chromosome 11 across all sequenced *Laverania* Genomes(Otto et al., 2018b). Assembly of this region is incomplete for the majority of strains. Plots begin 25kb before the rRNA loci on this region where the duplicated region between chromosomes 11 and 13 is. All assemblies of the strains that contain this region have this region shared between species and between chromosomes 11 and 13. The bottom x-axis shows the genomic distance from the starting point in each strain, but there is significantly more of the core genome to the left of the starting point.

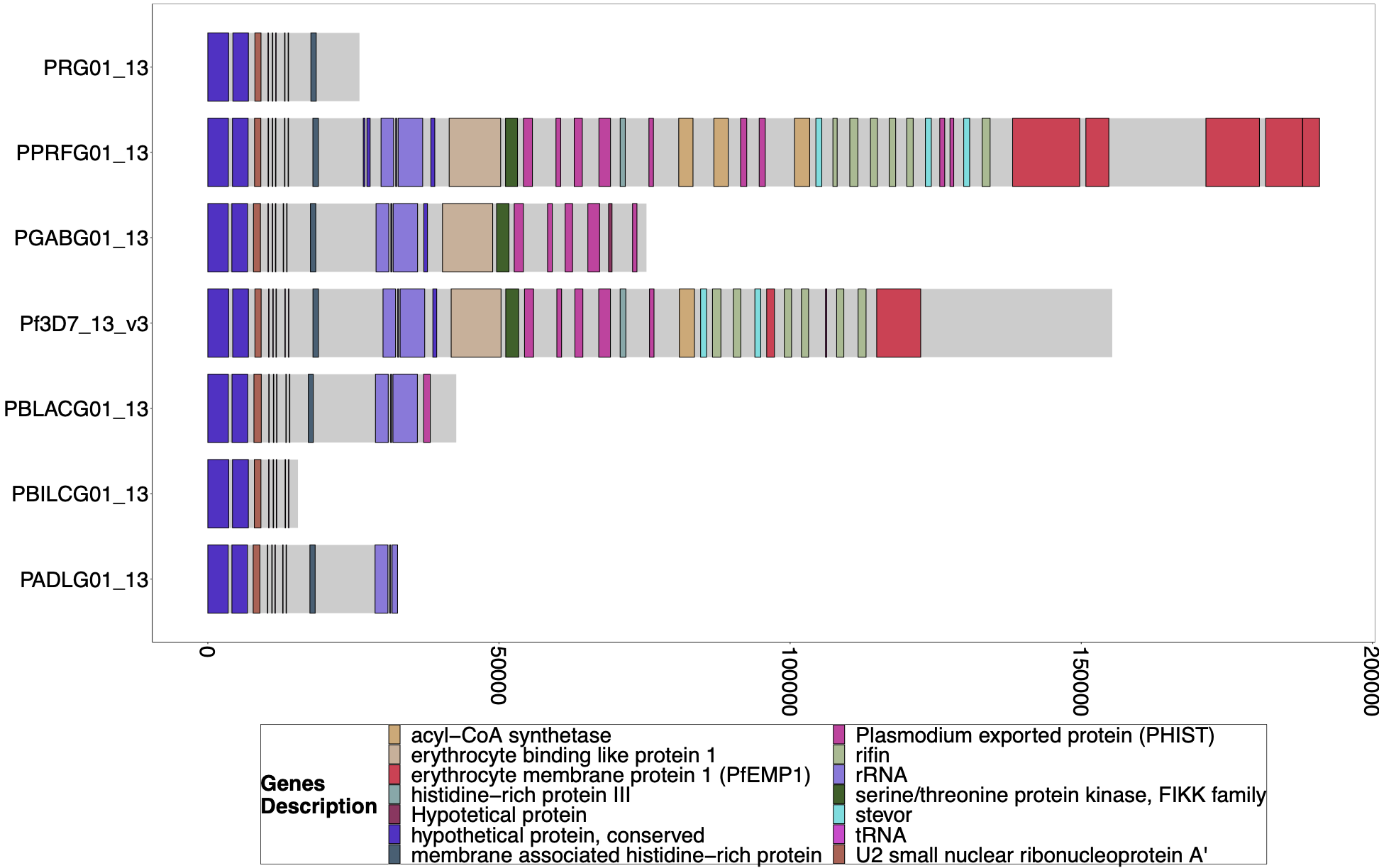

**Supplemental Figure 18****: Gene Annotations of Chromosome 13 of PacBio-assembled *P. Laverania* Genomes**

Plots of the peri-telomere regions of chromosome 13 across all sequenced *Laverania* Genomes(Otto et al., 2018b). Assembly of this region is incomplete for the majority of strains. Plots begin 25kb before the rRNA loci on this region where the duplicated region between chromosomes 11 and 13 is. All strains assemblies that contain this region have this region shared between species and between chromosomes 11 and 13. The bottom x-axis shows the genomic distance from the starting point in each strain, but there is significantly more of the core genome to the left of the starting point.

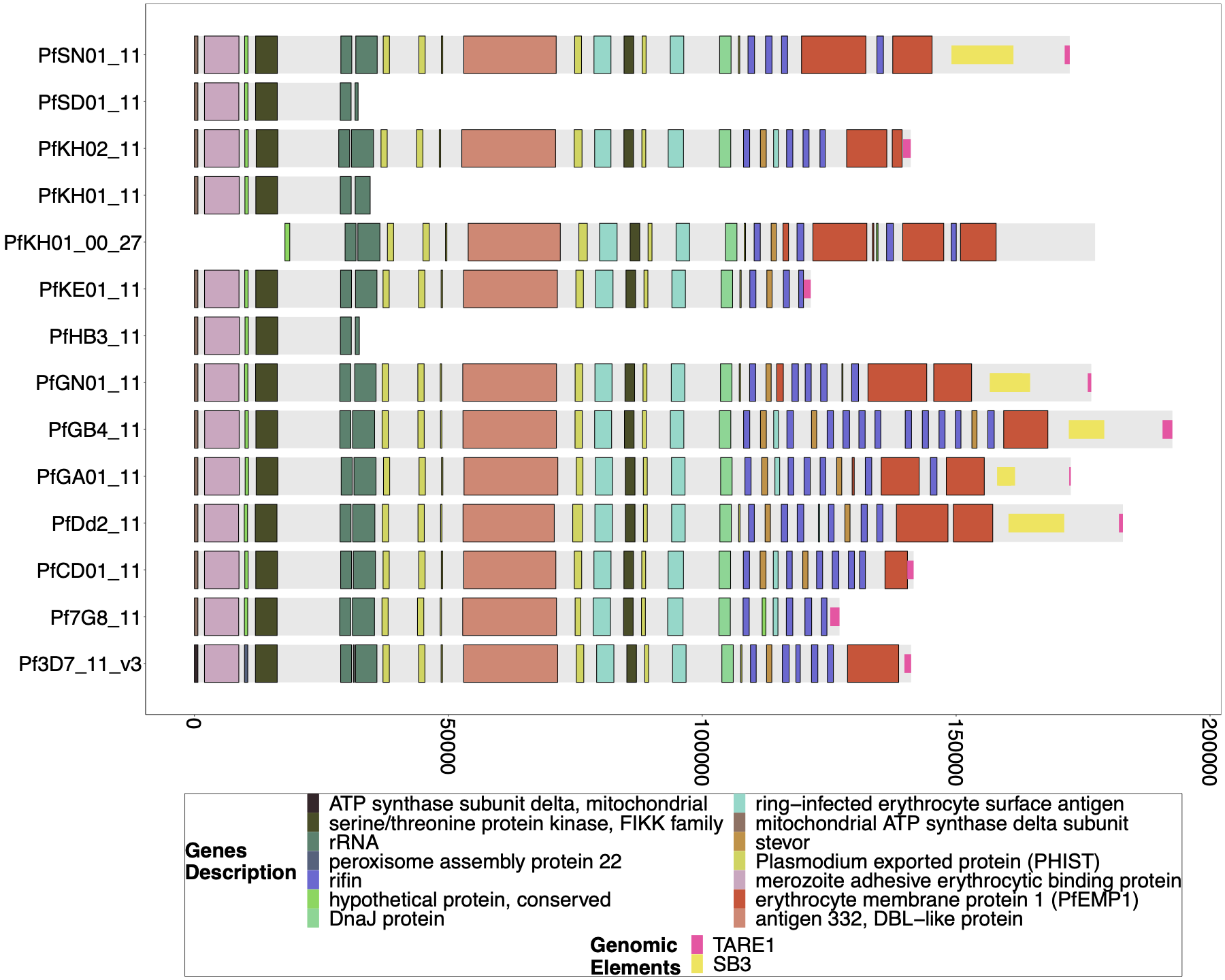

**Supplemental Figure 19****: Gene Annotations of Chromosome 11 of PacBio-assembled Genomes.** The genomic annotations across the 3` telomeric regions of PacBio-assembled genomes(Otto et al., 2018a) across chromosome 11 with the telomere repetitive elements (TAREs) are also shown if present. The presence of TAREs suggests that the assembly has made its way to the chromosome's sub-telomeric region (end). The previously published PacBio assembled genomes for SD01 and HB3 did not reach the TAREs for chromosome 11 and terminated in the segmental duplication. The absence of an assembled sub-telomeric region on chromosome 11 prevents a detailed analysis of the mechanism behind the deletion of *pfhrp3.* It is likely a result of the inability of the assembler and/or underlying PacBio reads to unambiguously traverse the segmental duplication and separate the duplicated chromosome 11 subtelomeric region sequence into two copies*.* The bottom x-axis shows the genomic distance from the starting point in each strain, but there is significantly more of the core genome to the left of the starting point.

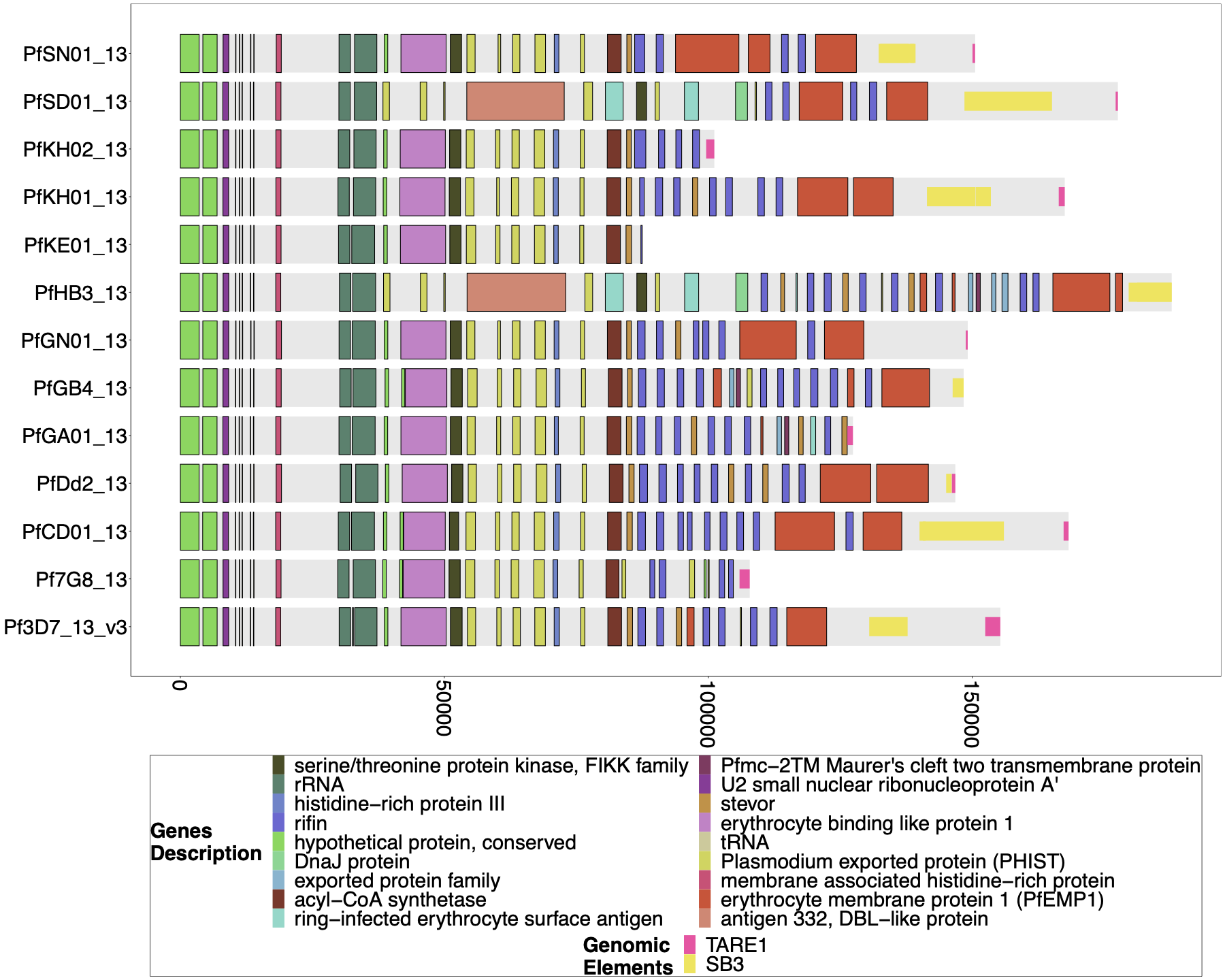

**Supplemental Figure 20****: Gene Annotations of Chromosome 13 of PacBio-assembled Genomes.** The genomic annotations across the 3` telomeric regions of PacBio-assembled genomes(Otto et al., 2018a) across chromosome 13 with the telomere repetitive elements (TAREs) are also shown if present. The presence of TAREs would suggest that the assembly has made its way all the way through the sub-telomeric region for the chromosome. The previously published PacBio-assembled genomes for SD01 and HB3 have sub-telomeric chromosome 11 sequences beginning after the segmental duplication, which suggests a translocation. Still, given the incompleteness of the chromosome 11 assembly in **Supplemental Figure 19**, it cannot be determined if this is simply a misassembly or a true translocation. The bottom x-axis shows the genomic distance from the starting point in each strain, but there is significantly more of the core genome to the left of the starting point.

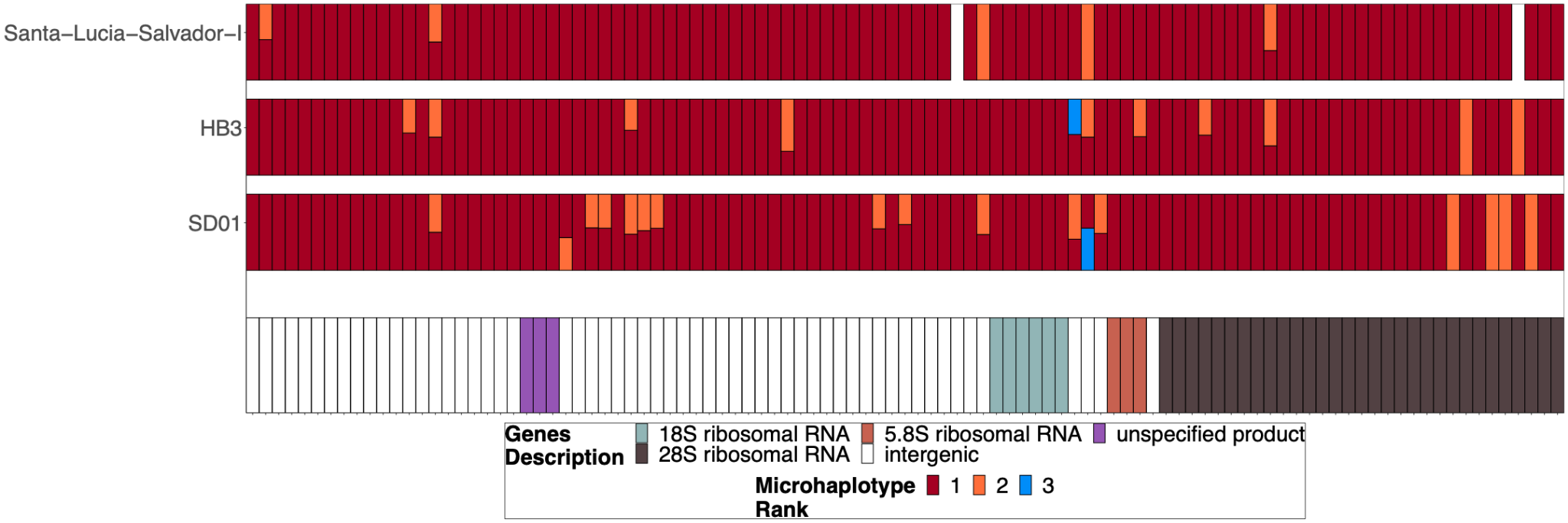

**Supplemental Figure 21****: Chromosome 11/13 15.2kb duplicated region for parasites SD01, HB3, and Salvador 1.** Subset of the parasites from **Supplemental Figure 15** but for SD01 and HB3, which were sequenced in this paper, and for Santa-Luca-Salvador-1, another lab isolate that shows similar *pfhrp3* deletion pattern 13^-^11^++^. SD01 and Santa-Luca-Salvador-1 have perfect copies, but Santa-Luca-Salvador-1 has variation at 3 loci within the duplicated region, and SD01 has variation at 13 loci within this region. HB3 has divergent copies of the duplicated chromosome 11 segment and contains variation within this region.

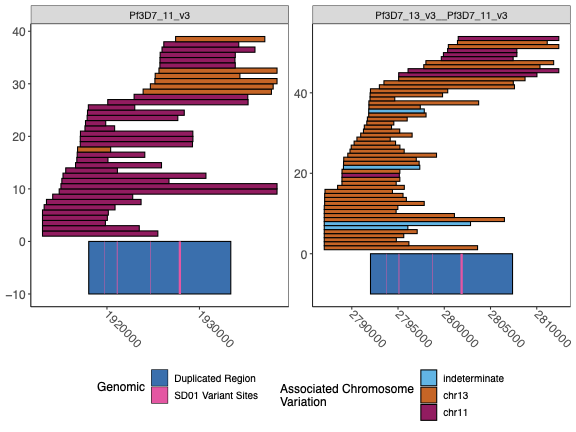

**Supplemental Figure 22****: Spanning PacBio and Nanopore Reads across the duplicated region for SD01.** The spanning Nanopore and PacBio reads across chromosomes chr11 and chr13 duplicated regions for isolate SD01. The visualization truncates the reads if they span outside of the range shown. The left panel is chromosome 11, and the right panel is the hybrid chromosome 13-11. The chr11/13 duplicated region is colored in dark blue on the bottom of the plot, and the 4 loci where the isolate SD01 has key variation within this region, which can be used to optimize bridging across this duplicated region, are colored pink. The reads are colored by the chromosome associated with the variation seen in each read. The association was made by linking the variation found within each of the 4 loci and looking at the reads spanning from each chromosome to see which variants were associated with which chromosome. Each locus had 2 variants and had a strong association with each chromosome.

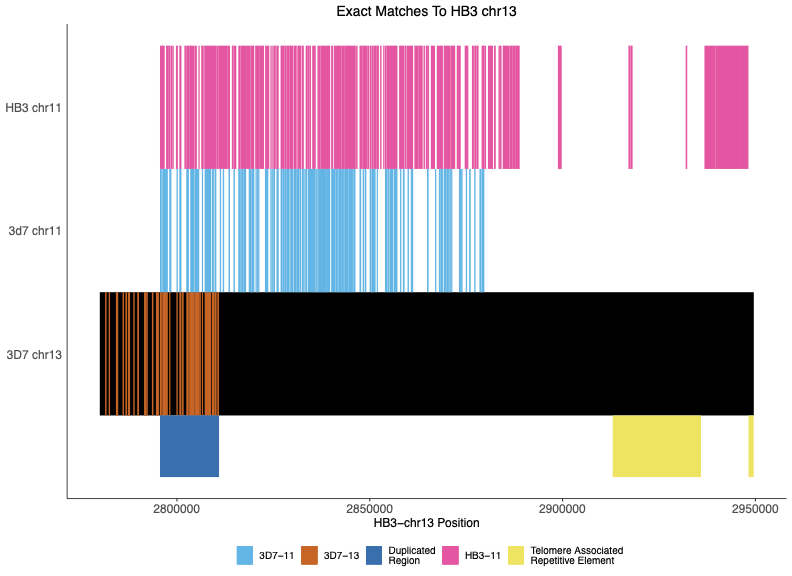

**Supplemental Figure 23****: Exact Matches between Nanopore-assembled HB3 chromosome 13 with HB3 chromosome 11, 3D7 chromosomes 11, 13.** The locations of exact matches between the Nanopore-assembled HB3 chromosome 13 and between the assembled chromosome 11 and the 3D7 11 and 13 chromosomes. The dark blue shaded region shows the location of the duplicated region between chromosomes 11 and 13. The assembled chromosome 13 matches the 3D7 chromosome 13 until this duplicated region and then more closely matches 3D7 chromosome 11 and its own chromosome 11. The figure begins at 50,000bp before the duplicated region, but the new HB3 chromosome 11 matches 3D7 chromosome 11 for the rest of the beginning of the contig.

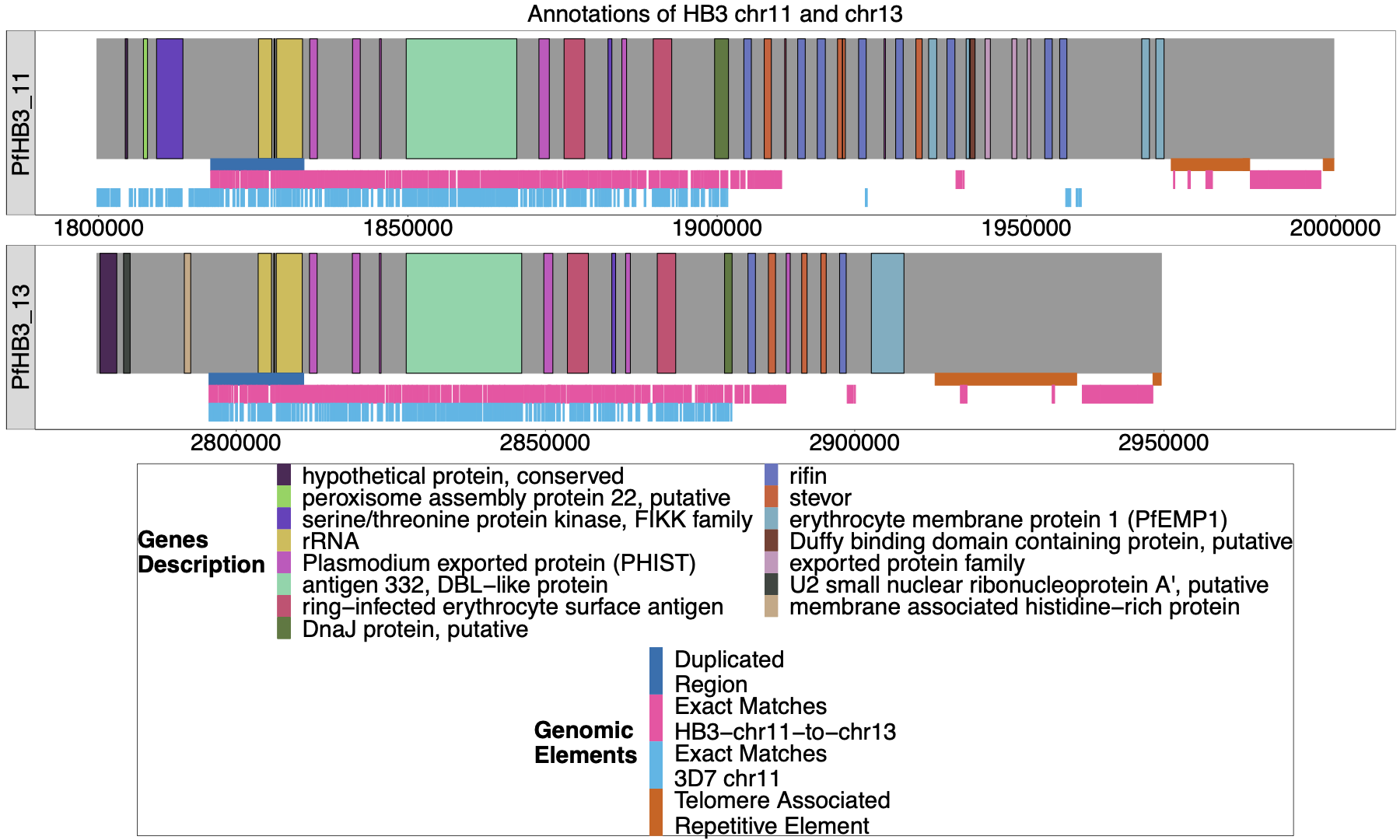

**Supplemental Figure 24****: Annotation of HB3 chromosomes 11 and 13-11.** The new Nanopore assembly of HB3 was annotated by Companion(Steinbiss et al., 2016), and the ends of chromosomes 11 and 13 are shown above. The duplicated region between chromosomes 11 and 13-11 is shown in blue under each chromosome, and the areas where HB3 chromosomes 11 and 13 have exact matches of at least 31bp are labeled in red underneath. Exact matches of at least 31bp to 3D7 chromosome 11 are shown in light blue. Both chromosomes end with telomere-associated repetitive elements (TARE), and both end with TARE1, which indicates that both assembled chromosomes reached the end of the telomere.

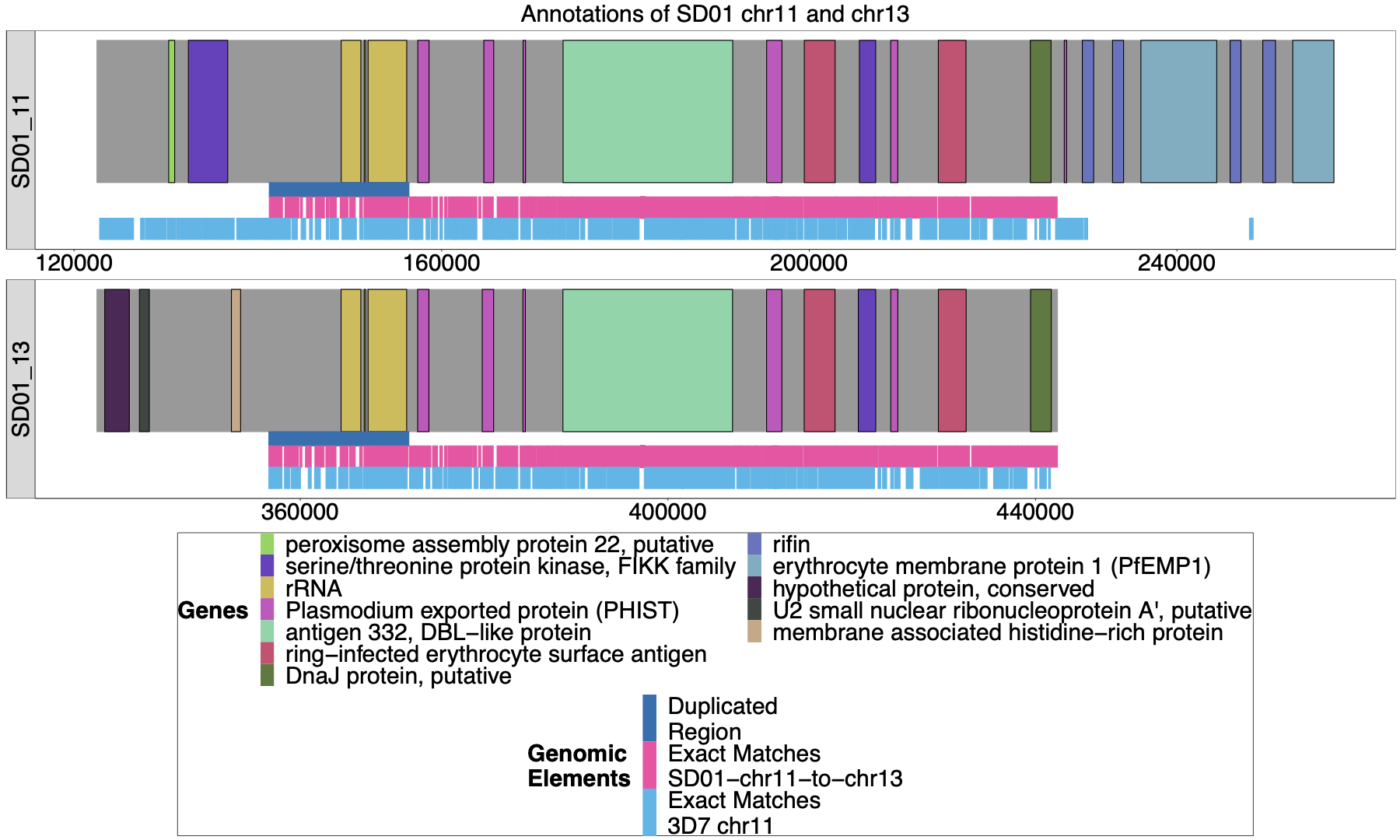

**Supplemental Figure 25****: Annotation of SD01 chromosomes 11 and 13.** The Nanopore assembly of SD01 was annotated by Companion(Steinbiss et al., 2016), and the ends of chromosomes 11 and 13 are shown above. The duplicated region between chromosomes 11 and 13 is shown in dark blue, and the areas where SD01 chromosomes 11 and 13 have exact matches of at least 31bp are marked out in pink underneath. Exact matches of at least 31bp to 3D7 chromosome 11 are shown in light blue. Due to the low quality of the input DNA of the SD01 parasite, the assembly of these chromosomes did not reach the end of the telomere, given that these assembled contigs did not contain TARE. The assembly of these two chromosomes shows a high degree of similarity from the duplicated region to the end of the 13 associated contig (98.4% similarity with only 1,428 difference over the 89,733 base region).

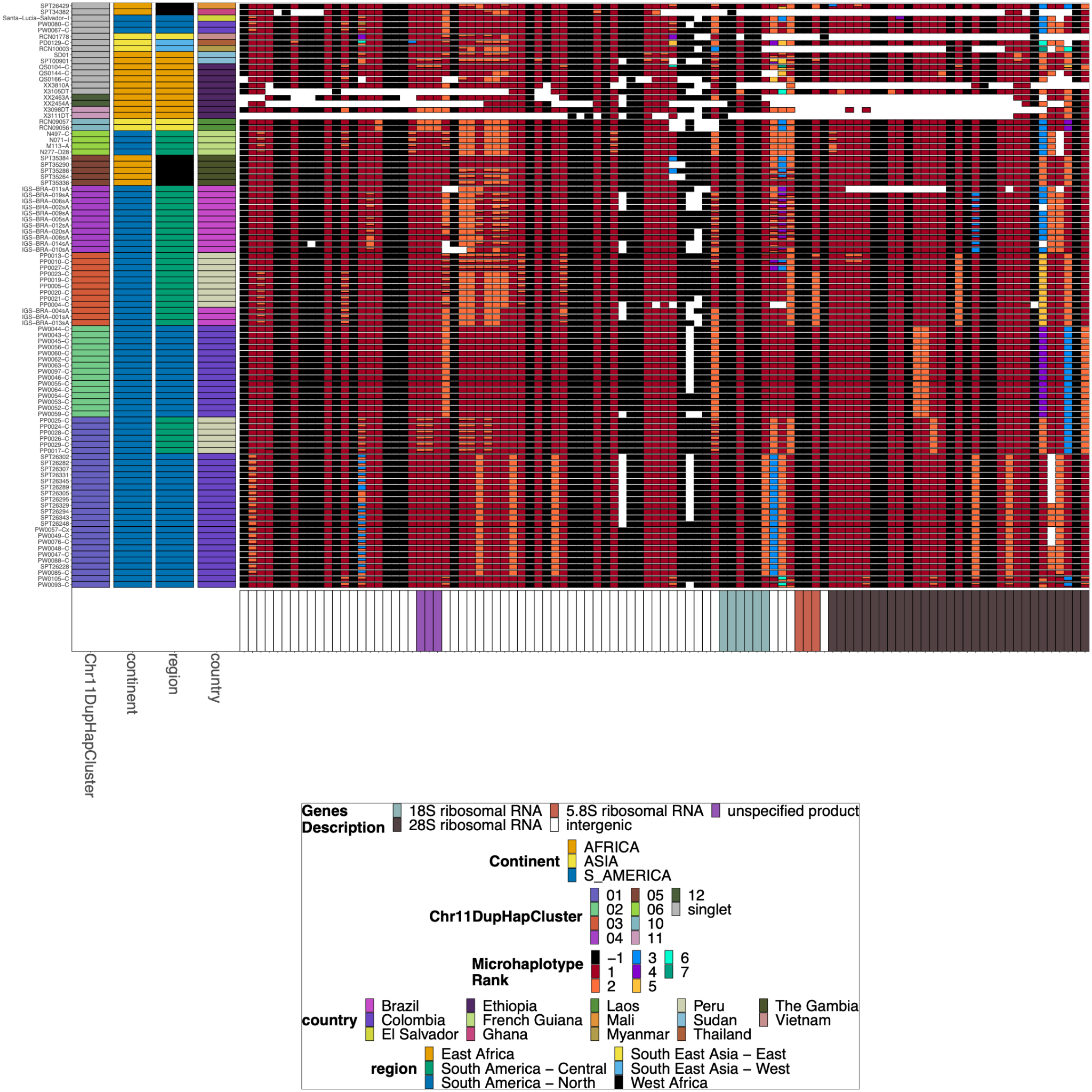

**Supplemental Figure 26****: Chromosome 11/13 15.2kb duplicated region for *pfhrp3* deletion pattern 13^-^11^++^ parasites with identical chromosome 11 segment haplotypes.**

A subset of parasites from **Supplemental Figure 15** for the chromosome 11/13 duplicated region for the parasites with identical chromosome 11 segments based on their microhaplotypes. The leftmost column contains the groupings based on the microhaplotypes on chromosome 11. Several parasites have divergent copies of the 15.2kb duplicated region despite the downstream chromosome 11 segments being a perfect copy. This would be consistent with the breakpoint for the duplication event within this region, where recombination occurred between nonidentical copies. The parasites are organized by the downstream duplicated chromosome 11 segment. For the 01, 03, 04, and 05 clusters, there are clear sub-clusters of haplotypes within this region consistent with separate duplication events creating the same downstream duplicated segment.

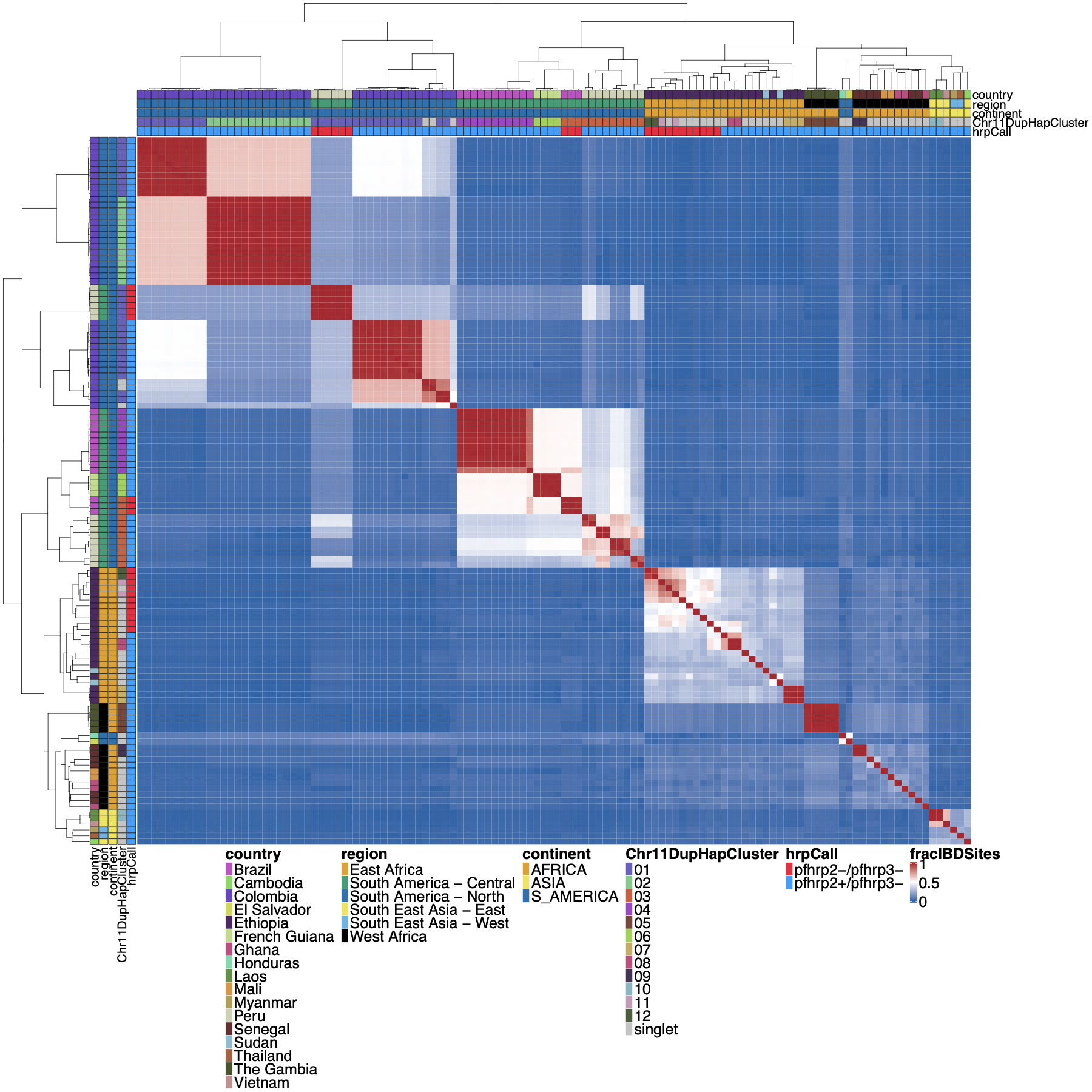

**Supplemental Figure 27**: **Whole genome** **IBD between** ***pfhrp3* deletion pattern 13^-^11^++^ parasites**

Whole genome identity by descent (IBD) was calculated between *pfhrp3* deletion pattern 13^-^11^++^ parasites using hmmIBD using whole genome biallelic SNPs. The fraction of sites in IBD between parasites was plotted above. The top and left side annotations are the same and contain the parasite's country, region, and continent of origin. The annotation also has the deletion calls for *pfhrp2/3* and which chromosome 11 haplotype cluster the parasite belongs to. Parasites clustered per continent of origin as expected, but there are different haplotype clusters per chromosome 11 duplicated clusters. When comparing the whole genome fraction of IBD sites, there are distinctive haplotype groups within the chromosome 11 haplotype cluster within the biggest clusters 01 (n=28) and 03 (n=12). The 01 cluster has 3 distinct groups, 03 cluster has 5 distinct groups. These groups are consistent with the distinct differences in the 15.2kb duplicated region (the region with the breakpoint for the duplication) (see **Supplemental Figure 26**), which would be consistent with different translocation events creating the same duplication segment of chromosome 11.

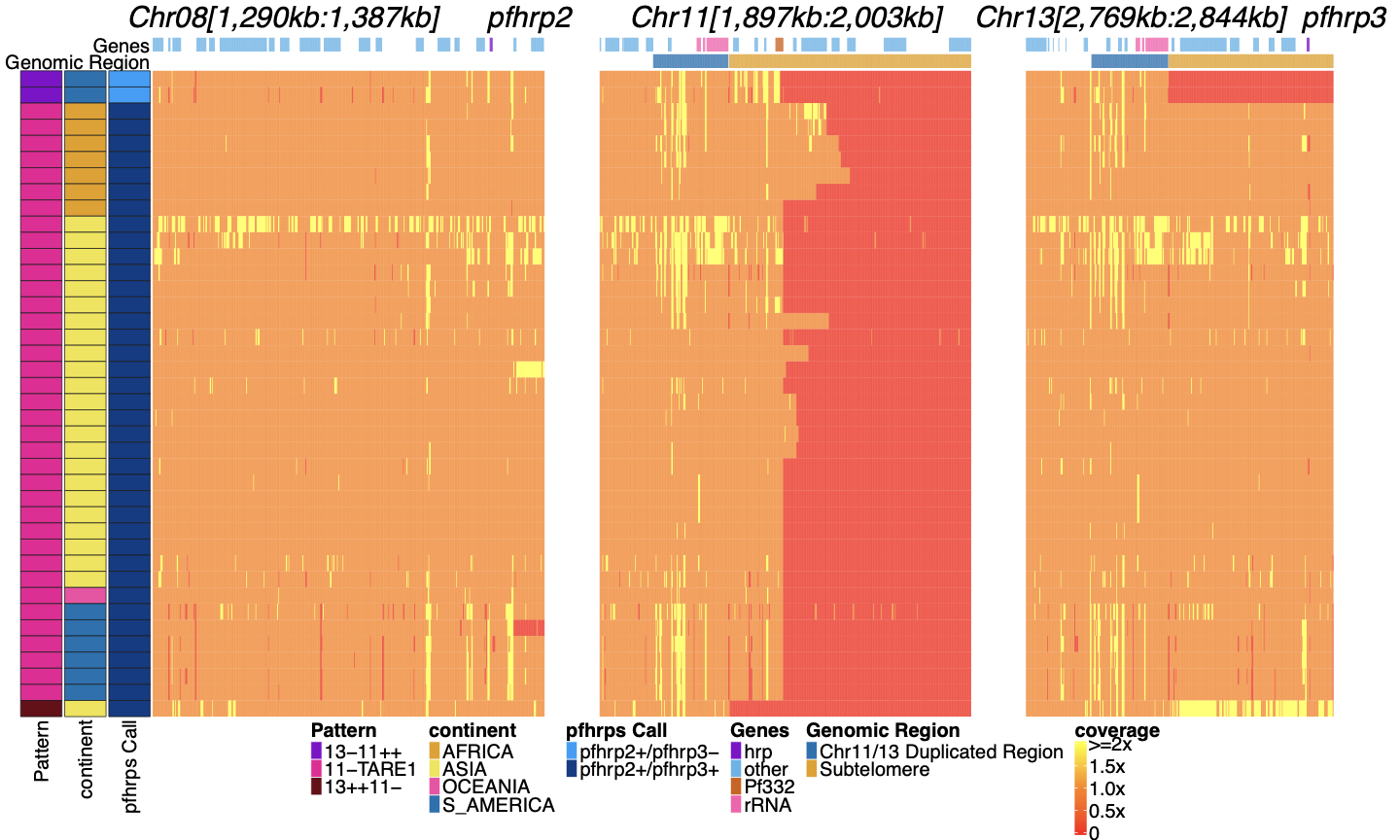

**Supplemental Figure 28****: Genome coverage chromosome 8, 11, and 13 regions of isolates with subtelomere deletion of chromosome 11.** Sequence coverage heatmap showing involved regions of chromosomes 11 (1,897,151 - 2,003,328 bp), 13 (2,769,916 - 2,844,785 bp), and 5 (944,389 - 988,747 bp) in the subset of the 19,313 parasites along with key lab isolates showing evidence of deletion of putative chromosome 11 subtelomere deletions. There are 41 parasites with evidence of chromosome 11 core genomic deletions. Each row is a parasite. The top annotation along chromosomes depicts the location of genes, and the second row delineates the duplicated region (dark blue) and subtelomere region (orange). The left parasite annotation includes the deletion pattern, continent of origin, and *pfhrp2/3* deletion calls. There were 41 parasites with evidence of sub-telomeric chromosome 11 deletions, 38 of which contained TARE1 sequence where coverage drops to zero, which would be consistent with telomere healing. Only one parasite (lab isolate FCR3) had deletion up and through *pf332* to the ribosomal duplicated region with subsequent duplication of chromosome 13 that would be consistent with the reciprocal of 13^-^11^++^. No field parasites had this pattern. The related clone of FCR3, IT, did not contain this pattern and would suggest that FCR3 duplicated this segment of chromosome 13 via translocation within culture and not in the field. The top 2 parasites show evidence of both deleting chromosome 13 with duplication of chromosome 11 consistent with the 13^-^11^++^ pattern and have deleted a portion of the chromosome 11 genome with evidence of TARE1 healing on chromosome 11. An unusual pattern that is only observed for these 2 parasites and not for the other 13^-^11^++^ pattern parasites.

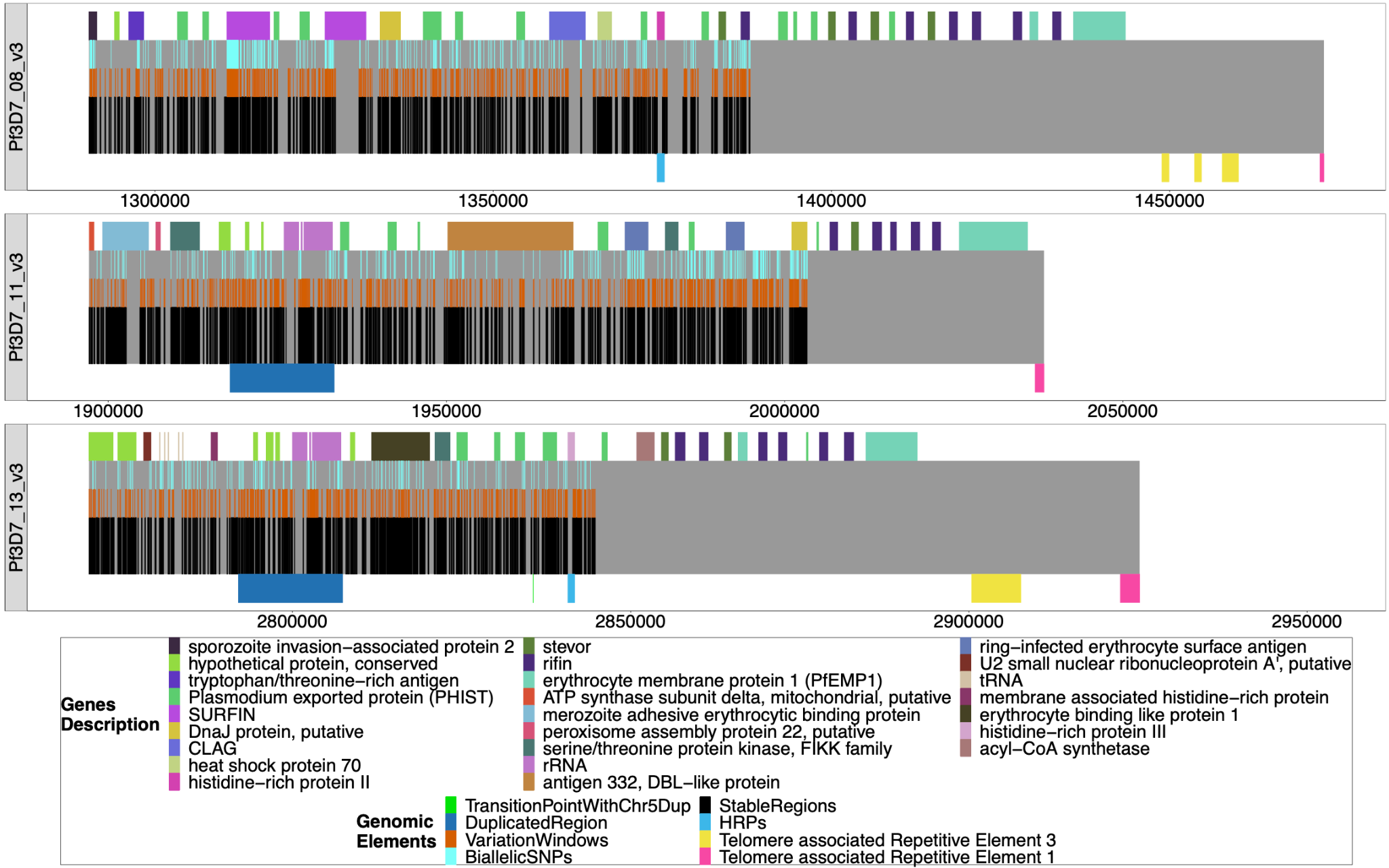

**Supplemental Figure 29****: Windows of interest chromosomes 8, 11, 13**. The chromosomes are mapped from the beginning of the regions of interest to the chromosomes’ ends, with all genes/pseudogenes annotations shown as colored on top of the gray bars representing the chromosomes. From top to bottom, the regions are from 3D7 chromosomes 8 (1290239-1387982), 11 (1897151-2003328), and 13 (2769916-2844785), and each span to the end of their chromosome. The black bars on the bottom half of each chromosome are non-paralogous regions present in all strains, as described in the Methods section. The last black bar to the end of the gray bar represents the sub-telomeric regions that are not homologous between strains. The orange bars on top of the black bars are sub-regions where there is variation that can be used to type the chromosomes.The light blue bars on top of the organe bars are the presence of biallelic SNPs. The duplicated region between chromosomes 11 and 13 is shown (dark blue bars below chromosomes 11 and 13), as are the regions containing the *pfhrp* genes (lighter blue bars below chromosomes 8 and 13). The yellow (TARE3) and pink (TARE1) bars on the bottom of the chromosomes represent the telomere-associated repetitive elements found at the end of chromosomes.

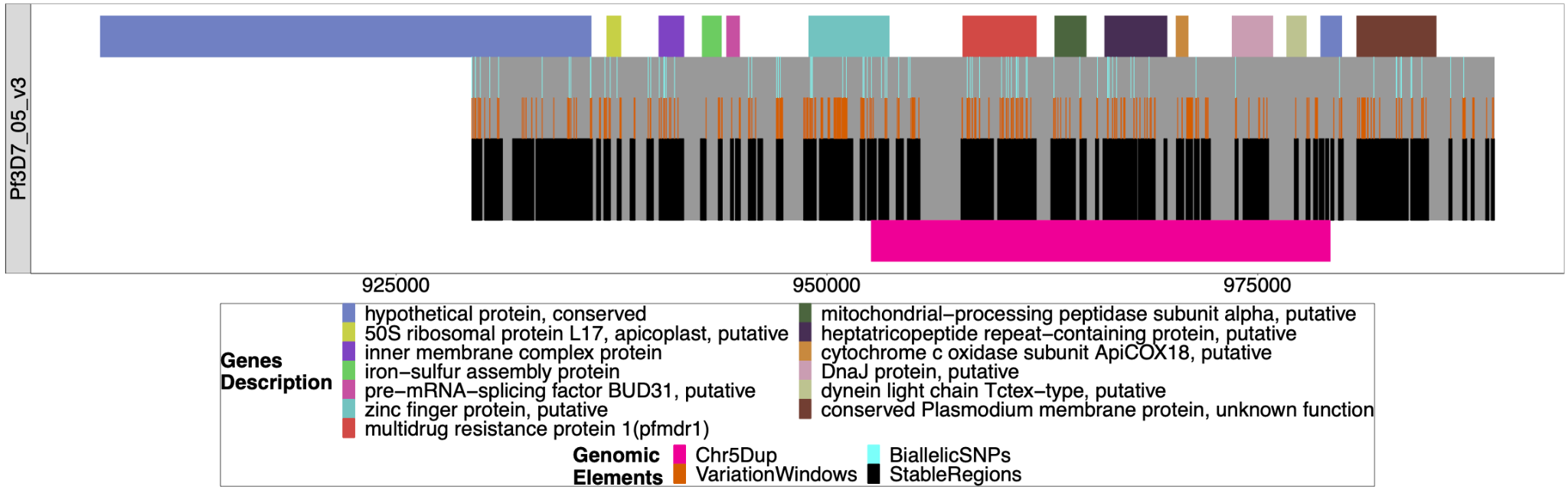

**Supplemental Figure 30****: Windows of interest chromosome 05 around *pfmdr1***

The windows used to investigate the duplication around *pfmdr1* on chromosome 5 associated with the deletion of *pfhrp3*. All genes/pseudogenes annotations are shown on top of the gray bars representing the chromosome region investigated (929384-988747). The first gene spans outside of this region and was shown to overhang this region, but the content underneath was not investigated. The black bars on the bottom half of each chromosome are non-paralogous regions present in all strains, as described in the Methods section. The orange bars on top of the black bars are sub-regions with variation that can be used to type the chromosome. The light blue bars on top of the organe bars are the presence of biallelic SNPs. The pink bar shows the region that is duplicated in pattern 13^-^5^++^.

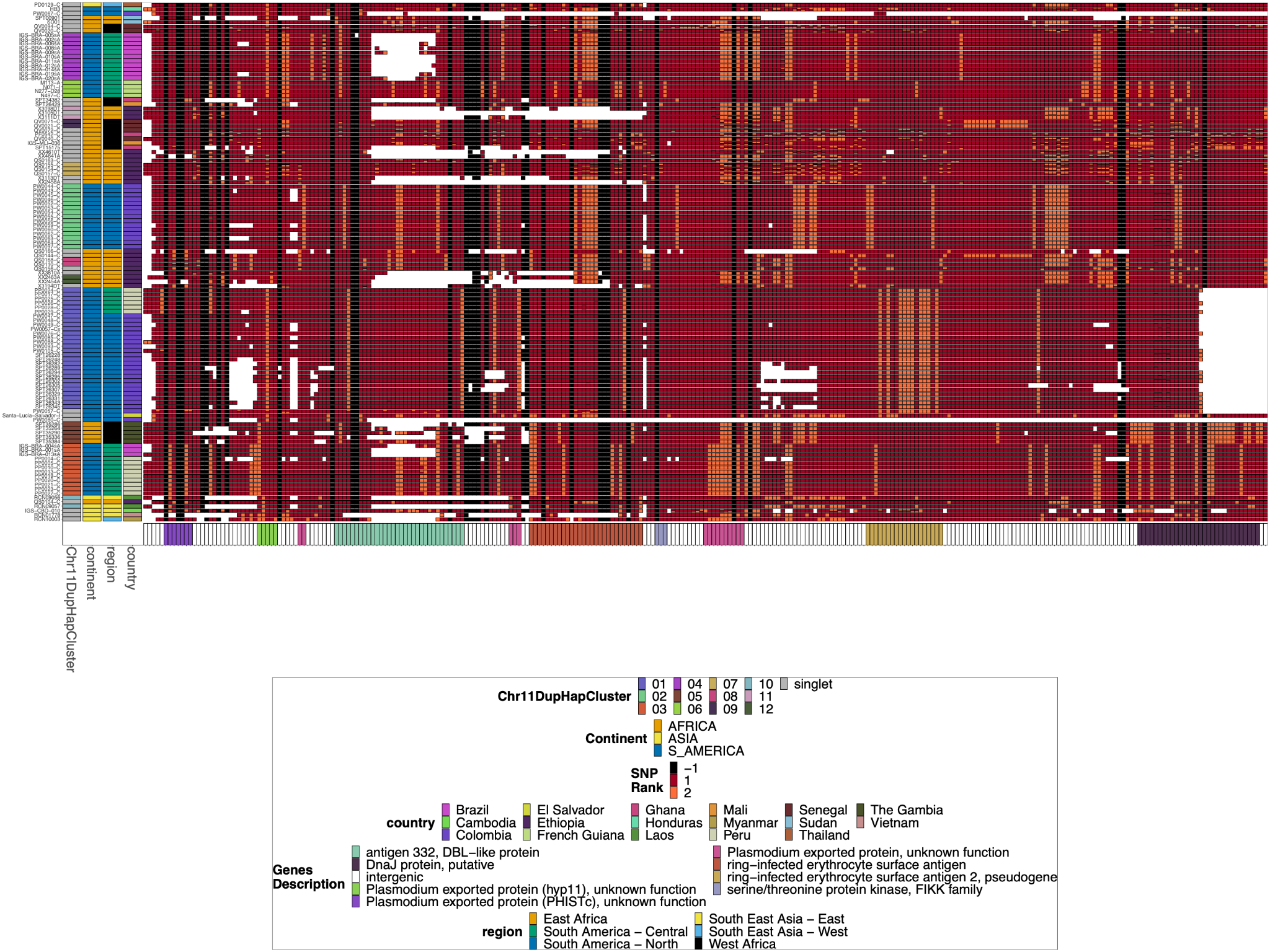

**Supplemental Figure 31**: **Chromosome 11 Duplicated Segment *pfhrp3* deletion Pattern 13^-^11^++^ parasites Biallelic SNPs**

This plot was created and annotated the same way as **Supplemental Figure 11**, but biallelic SNPs were used instead of small windows of microhaplotypes. The Chr11DupHapCluster is the same grouping as determined in **Supplemental Figure 11**, and these clusters group together similarly when clustered by the biallelic SNPs.

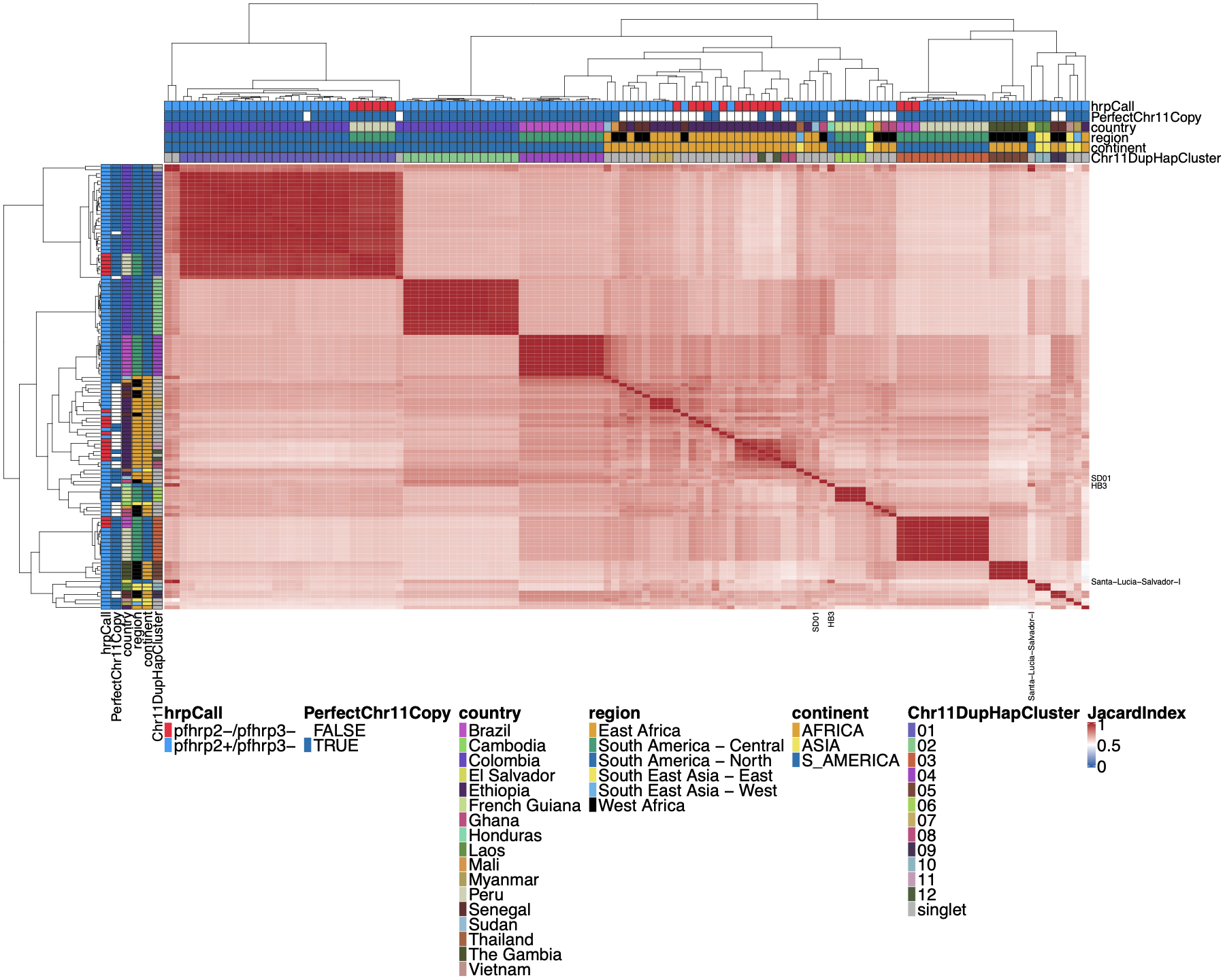

**Supplemental Figure 32**: **Jaccard similarity using Biallelic SNPs between parasites for chromosome 11 duplicated segment for *pfhrp3* deletion pattern 13^-^11^++^ parasites**

This plot was created and annotated in the same way as **Supplemental Figure 10**, but biallelic SNPs were used instead of small windows of microhaplotypes. The Chr11DupHapCluster is the same grouping as determined in **Supplemental Figure 10**. Using biallelic SNPs created clustering that was very similar to that based on the microhaplotype windows.
